# Surface Complex V couples proton gradient across the plasma membrane to ATP production in cancer

**DOI:** 10.64898/2026.05.19.726308

**Authors:** Samantha A. McLaughlin, Kendall W. Knechtel, Zelia M. Correa, J. William Harbour, Daniel Pelaez

**Author notes:** **Corresponding Author:** Daniel Pelaez, PhD, Associate Professor of Ophthalmology, Spencer Chair in Retinal Research, 1638 NW 10^th^ Ave. Miami, FL 33178.

## Abstract

Cancer cells alter their metabolism to support growth and survival, most notably by fermenting glucose to lactate even in the presence of oxygen, a phenomenon known as the Warburg effect. Although this metabolic state has been recognized for decades, its bioenergetic advantages remain unclear, as fermentation produces less net ATP than mitochondrial respiration. How aerobic fermentation contributes to cellular energy balance therefore remains unresolved. Here, we show that extracellular acidification generated by lactate export creates a proton gradient across the plasma membrane that is harnessed by ectopic ATP synthases to drive intracellular ATP production. We find that ATP synthase and proton-shuttling components of the mitochondrial respiratory chain translocate to the plasma membrane in cancer cells and are preferentially oriented to exploit this gradient, linking a hallmark of aerobic fermentation directly to energy supplementation. This work provides a mechanistic resolution to the apparent energetic inefficiency of the Warburg paradigm and identifies a previously unrecognized pathway for energy complementation in cancer.

## Introduction

Cancer cells undergo profound metabolic reprogramming to sustain proliferation and survival, most notably through the preferential conversion of glucose to lactate even in the presence of oxygen and fully functional mitochondria, a phenomenon known as the Warburg effect. Although this metabolic phenotype has been recognized for nearly a century, the functional advantage conferred by aerobic glycolysis and fermentation remains incompletely understood. The defining features of the Warburg phenotype, namely high glycolytic flux, sustained lactate export, and extracellular acidification, are consistently selected for during tumor evolution, but how they fit into the cellular energetic framework, given the apparent energetic inefficiency of glycolysis, has yet to be fully elucidated.

Multiple non-mutually exclusive hypotheses have been proposed to explain the Warburg effect, including compensation for slower oxidative phosphorylation kinetics^1–7^, regeneration of NAD^+^ to support anabolic metabolism^1,8–10^, and saturation of mitochondrial respiration capacity^11–13^. Additional models suggest roles in limiting reactive oxygen species^14,15^, enabling adaptation to hypoxia^16^ or nutrient stress^2,17–23^, or promoting tumor progression through microenvironmental acidification^24,25^. While these models explain specific aspects of cancer biology, none provides a mechanistic explanation for how sustained lactate export and extracellular acidification are coupled to net cellular energetic yields, leaving the teleological advantage (evolutionary and mechanistic basis) of aerobic glycolysis unresolved.

Here, we uncover that a missing mechanistic link underlying the Warburg effect is the presence of functional ATP synthase complexes at the plasma membrane. Lactate-driven extracellular acidification generates a proton gradient across the plasma membrane, analogous to the proton motive force across the inner mitochondrial membrane, which can be harnessed for ATP synthesis outside of mitochondria. Although ATP synthase is canonically localized to the inner mitochondrial membrane, accumulating evidence indicates that it can translocate to the cell surface as ectopic ATP synthase (ecto-ATP synthase) ^26–35^, where its abundance is correlated to tumor aggressiveness^32,33,36,37^, although its functional role in this context remains unclear.

We demonstrate that components of the mitochondrial respiratory chain are embedded within the plasma membrane in retinoblastoma cells, where they are dynamically regulated by glycolytic demands and retain catalytic function. Under physiologically relevant conditions, ATP synthase is preferentially oriented with its proton-shuttling F_O_ domain facing the extracellular space, enabling it to exploit the proton gradient generated by lactate export to drive intracellular ATP synthesis. This establishes a direct mechanistic link between aerobic glycolysis and fermentation to cellular energy production, thereby reconciling a net-positive energetic architecture for the Warburg effect. Finally, we advance the proton pore of the extracellular F_O_ domain as a selective and targetable metabolic vulnerability, providing a potential therapeutic strategy to disrupt cancer bioenergetics.

## Methods

### Cell Culture

Patient-derived Rb cell lines (RB006, RB036) were derived from the tumors of enucleated eyes (University of Miami Institutional Review Board [no. 20130636]) and cultured at 5% O_2_ in Dulbecco’s Modified Eagle Medium/Nutrient Mixture F-12 with L-glutamine (Gibco) with 2% B-27 supplement (50X) without vitamin A (Gibco), 1% antibiotic-antimycotic (100X) (Gibco), 1% MEM non-essential amino acids (100X) (Gibco), human basic fibroblast growth factor (10 ng/ml; PeproTech, catalog no. 100-18B), recombinant human stem cell factor (10 ng/ml; PeproTech, catalog no. 300-07), and recombinant human epidermal growth factor (20 ng/ml; PeproTech, catalog no. AF-100-15). Human MCF-7 cell lines were obtained from the American Type Culture Collection (ATCC) and maintained in DMEM supplemented with 10% heat-inactivated fetal bovine serum (FBS) (Thomas Scientific) and 1% antibiotic-antimycotic. Dental pulp-derived mesenchymal stem cells were obtained from the ATCC and cultured in DMEM supplemented with 10% heat-inactivated FBS, 1% antibiotic-antimycotic, and 2mM glutaMAX (Gibco). HEK293 were obtained from the ATCC and cultured in DMEM supplemented with 10% heat-inactivated FBS, 1% antibiotic-antimycotic, and 1% MEM non-essential amino acids (100X) (Gibco) Cell proliferation and viability were measured using Trypan Blue Stain (Gibco). All cells were cultured at ambient oxygen unless otherwise stated. All hypoxic experiment were performed under 1% oxygen, 5% carbon dioxide with nitrogen balance (Airgas, Part # X03NI94C2000650)

### Plasmids and lentivirus expression vectors

The cell lines listed below were established by stable transduction. Adherent cell lines (HEK293, MSC) were plated in 10 cm^2^ plates and grown to 90% confluency prior to the addition of approximately 9 × 10^8^ TU/ml of recombinant lentivirus in fresh culture media. After 48 hours of incubation, the virus was removed, and the cells were rested in normal culture media for 48 hours, followed by selection with 1ug/mL of puromycin. For suspension cell lines (RB006, RB036), approximately 3 × 10^8^ TU/ml of recombinant lentivirus was added to the Rb cells cultured in 6-well plates and centrifuged for 1 hour at 1000 x g at 32C. After 48 hours of incubation, cells were transferred to T75 flasks for recovery. After 48 hours, cells were selected with 1ug/mL of puromycin. EGAP expression was confirmed by fluorescence imaging.

| Cell Line | Gene/Fusion | Backbone | Promoter | Antibiotic Resistance | Source |
| --- | --- | --- | --- | --- | --- |
| RB036, HEK293 | ATP5F1B-EGFP | pLVX | EF1A | Puromycin | Vector Builder |
| RB006, RB036, MSC | ATP5ME-HiBiT | pLVX | EF1A | Puromycin | Vector Builder |
| RB006, RB036, MSC | ATP5F1B-HiBiT | pLVX | EF1A | Puromycin |  |
| RB006, RB036, HEK293 | ActA-mCherry | pLVX | EF1A | Puromycin | Addgene (#177433) |

pATP5F1B-HiBiT was engineered by removing the EGFP protein from the backbone pATP5F1B-EGFP (Vector Builder) and inserting the 11-amino acid HiBiT peptide (VSGWRLFKKIS). Briefly, purified pATP5F1B-EGFP was amplified with forward and reverse primers, followed by Dpn1 digestion and T4 DNA ligation. The resultant pATP5F1B-HiBiT was transformed into NEB 5-alpha competent E. coli (C2987H) and plated on a selection plate for incubation overnight at 37°C. Colonies were then inoculated into selection LB broth overnight at 37°C. Individual colonies were picked and sent for sequencing validation at Plasmidsaurus.

Forward primer: 5’-GTGTCTGGCTGGAGACTGTTCAAGAAGATCAGCTAAACCCAGCTTTCTT GTACAAAGTGGTGATAATCGA-3’

Reverse primer: 5’-[Phos] TGATCCGCCGCCACCCGA-3’

### Immunofluorescence

For live-cell immunostaining, cells were cultured with the indicated antibody in culture media for 1 hour at 37°C, washed thrice with FluoroBrite DMEM (Thermo Fisher) by centrifugation at 300 RPM for 3 min, and subsequently stained with CellMask Deep Red (Thermo Fisher) according to the manufacturers protocol. After washing, cells were resuspended in FluroBrite DMEM and imaged at 63X using an SP8 Leica laser scanning confocal microscope (Leica Microsystems Inc.). For fixed-cell immunostaining, cells were seeded on coverslips coated in poly-l-lysine for 24 hours. They were then fixed in 4% paraformaldehyde (PFA) in PBS at room temperature for 20 minutes and washed 3X with PBS. To examine surface expression of ATP synthase, the cells were blocked with 5% bovine serum albumin (BSA) for 1 hour at room temperature, and probed with the primary conjugated antibody overnight at 4°C. Coverslips were mounted onto the glass slides with SlowFade Diamond Antifade mounting medium (Thermo Fisher) and imaged using a SP8 Leica laser scanning confocal microscope.

For live-cell imaging of pATP5F1B-EGFP, pActA-mCherry, or dual-expressing cells, suspension cells (RB006, RB036) were cultured in 24-well plates and adherent cells (HEK293) were grown on poly-l-lysine coated coverslips for 24 hours. Cells were washed 1X with PBS, stained with CellMask Deep Red, washed 3X with FluroBrite DMEM, and imaged immediately on an SP8 Leica laser scanning confocal microscope. Colocalization was analyzed using Fiji/ImageJ and the Coloc2 plugin to calculate the Thresholded Manders’ Coefficients (tM1, tM2). Values ranged from 0 to 1.

### Plasma membrane isolation

Cells were cultured in T75 tissue culture flasks with culture media until 90% confluency, at which point flasks were either maintained at normal oxygen conditions or transferred to 1% O_2_ for 12 hours. Plasma membrane and mitochondrial fractions were isolated using the Minute™ Plasma Membrane/Protein Isolation and Cell Fractionation Kit (Invent Biotechnologies, Inc.) according to the manufacturers protocol. Isolated cellular fractions were resuspended in RIPA lysis and extraction buffer (Thermo Scientific) with Pierce protease and phosphatase inhibitor mini tablets (Thermo Scientific) and stored at −80°C until further analysis.

### Western blot

Cell pellets were resuspended in RIPA lysis buffer with protease and phosphatase inhibitors. Resulting cell lysates were incubated on ice for 30 min with intermittent vigorous vortexing for 30 s, centrifuged for 15 minutes at 20,000 x g to remove cellular debris, and quantified with QuBit Protein Broad Range Assay (Thermo Fischer). Equal protein quantities were denatured by boiling with SDS sample buffer for 10 minutes at 95°C with 300 RPM. Proteins (10µg per condition; 25µg per condition for plasma membrane isolation) were resolved on SDS-polyacrylamide gel electrophoresis gradient gels and transferred to a polyvinylidene difluoride membrane. Blots were blocked in 5% bovine serum albumin (BSA) in 0.15% Tween 20 in tris-buffered saline (TBS-T), probed for 1.5 hours at room temperature with indicated primary antibodies in 2.5% BSA/TBS-T, washed with TBS-T, and incubated for 1 hour with a secondary antibody in TBS-T. proteins were visualized using SuperSignal chemiluninsecence on iBright 1500 (Invitrogen) or ChemiDoc (BioRad).

### Mass spectrometry

Isolated plasma membrane pellets from RB006 and RB036 cells cultured in either normal oxygen (21% O_2_) or 1% O_2_ and immediately placed in dry ice. CoIP samples were shipped to the Proteomics and Metabolics Core Facility at H. Lee Moffitt Cancer Center and Research Institute (Tampa, FL) for liquid chromatography-tandem mass spectrometry (LC-MS/MS).

### Flow cytometry

Cells were washed with PBS once by centrifugation at 300 x g for 5 minutes at room temperature, strained using a 40um cell strainer and adjusted to a concentration of 1×106 cells/ml. The intact cells were incubated with the primary antibodies: Alexa Fluor® 555 Anti-ATP synthase C antibody [EPR13907] (Abcam, ab210732)) and Alexa Fluor® 647 Anti-ATPB antibody [3D5] (Abcam, ab197649) at 10ug/mL in FACS Buffer (PBS with 1% bovine serum albumin) at 4C for 1 hour. Cells were washed three times in FACS buffer by centrifugation. 1uL of LIVE/DEAD™ Fixable Near-IR Dead Cell Stain (Invitrogen) was added per 1mL of cells, vortexed, and incubated at room temperature for 30 minutes. Cells were washed via centrifugation twice in FACS Buffer, strained, and resuspended in 4% formaldehyde for 15 minutes at room temperature. Controls were either unstained, stained with a single antibody, or stained with LIVE/DEAD™ only. After centrifugation, cells were resuspended in FACS buffer and analyzed immediately on a FACSymphony A5 SE (BD Biosciences).

### HiBiT-LgBiT

Nano-Glo® HiBiT Extracellular Detection System (Promega) was used with cells stably expressing either ATP5F1B-HiBiT or ATP5ME-HiBiT. Cells were plated at a density of 1 × 10^4^ cells/well in a 96-well tissue culture treated plate and cultured for 12 hours under either normoxic (21% O_2_) or hypoxic (1% O_2_) conditions. For detection of cell surface expression, equal volume of nonlytic detection reagent containing the substrate furimazine and Large BiT. For detection of total HiBiT-tagged protein(intracellular and extracellular), Triton X-100 was to the detection reagent at a final well concentration of 0.05% (v/v). Non-transformed cells were used as background controls. The samples were mixed with gentle pipetting. Luminescence was recorded with a SpectraMax i3 (Molecular Devices) 10 minutes after adding reagent.

### Transmission electron microscopy

All steps were performed at room temperature with centrifugation after each wash for 2 minutes at 500 rpm. The cell pellet was rinsed briefly twice in DPBS followed by fixation for 1 hour in 2% paraformaldehyde. Cells were washed three times in DPBS with 50 mM glycine then fixed in 2.5% glutaraldehyde in 0.1M sodium cacodylate buffer with CaCl_2_ for 1 hour at room temperature. Samples were submitted to the transmission electron microscopy core for traditional electron microscopy processing. Imaging was done on JEM-1400 (JEOL Ltd.).

### Co-immunoprecipitation

Cells were cultured in normal culture media under either normoxic or hypoxic conditions for 12 hours. Cells were lysed with cold Pierce IP Lysis Buffer (2 5mM Tris-HCl [pH 7.4], 150 mM NaCl, 1 mM EDA, 1% NP-40, 5% glycerol) (Thermo Fischer) with protease and phosphatase inhibitors, incubated on ice for 30 minutes with periodic vigorous vortexing, and pelleted by centrifugation at 13,000 x g for 10 minutes. Protein was quantified by Qubit Protein Broad Range Assay (Invitrogen) and all samples were adjusted to a concentration of 1 mg/mL. Co-immunoprecipitaiton was performed using DynaGreen CaptureSelect Anti-IgG-Fc (Multi-Specia) Magnetic Beads (Invitrogen) according to the manufactuerers protocol. Briefly, 4µg of antibody diluted in PBS with 0.05% Tween 20 was incubated with the DynaGreen magnetic beads with rotation for 30 min at room temperature. Sample containing the antigen was then added to the magnetic bead-antibody complex and again incubating with rotation for 30 minutes at room temperature. For analysis by LC-MS/MS, after washing, magnetic beads with the bound antibody/protein complex were frozen at −80°C until analysis. For immunoblot, the target antigen was eluted with SDS sample buffer for 10 minutes at 70°C, then 25µl of supernatant was loaded onto a SDS-polyacrylamide gel electrophoresis gradient gel following the method under “Western blot”.

### Cell surface ATP generation assays

Adherent cells were detached with trypsin-EDTA (0.25%) for 5 min at room temperature followed by inactivation, centrifugation at 100 rpm for 5 min, and resuspension in normal media. Suspension cells were collected by centrifugation. All cells were seeded at 800,000 cells/well in 6-well plates. For acidification experiments, after 24 hours of incubation, cells were treated with media that had been acidified to the indicated pH using 1N HCl. Cells were cultured in media supplemented with 5mM HEPES and incubated in a non-CO_2_ incubator. At each time point, cells were collected by centrifugation at 300 rpm for 5 min. The supernatant was collected and the pH measured using sension+ PH31 (Hach). The extracellular fraction was also assayed for ATP production by Cell-Titer Glo 2.0 luminescence assay (Promega). The remaining intracellular fraction was resuspended in DMEM and also assayed for ATP production with Cell-Titer Glo 2.0. Luminescence was recorded with a SpectraMax i3 (Molecular Devices).

For alkalinization assays, cells were collected and plated as above and then treated with 20nM of AZD3965, a specific MCT1 inhibitor (MedChemExpress). For rescue experiments, the media was re-acidified with HCl (5mM). Extracellular pH and extracellular and intracellular ATP production were assayed as above.

### Cell surface MRC complex inhibition

Cells were treated with either Rotenone/Antimycin A (0.125 µM), Oligomycin A (0.375 µM), or carbonyl cyanide-p-trifluoromethoxyphenylhydrazone (FCCP) (0.125 µM) and cultured in media supplemented with 5mM HEPES and incubated in a non-CO_2_ incubator. Media pH was measured every hour using sension+ PH31 (Hach). Control cells were cultured under either experimental conditions (no drug) or with normal media in a CO_2_ incubator. Control and experimental conditions were compared by measuring media pH and cell viability with Trypan Blue exclusion assay at each time point.

### Bu_2_Sn(of)Cl synthesis

#### Dose-response curves

Cells were seeded at a density of 1 × 10^4^ cells/well in a 96-well tissue cultured treated plate and treated with Bu_2_Sn(of)Cl (64 – 0.0001µM) for 72 hours under normal culture conditions. Dimethyl sulfoxide (DMSO) was used as vehicle control. Cell viability was measured with either an MTT 3-(4,5-dimethylthiazol-2-yl)-2,5-diphenyltetrazolium bromide cell viability assay (Biotium) or CellTiter-Glo 2.0 cell viability assay (Promega) according to the manufacturers protocol.

## Statistical Analysis

All statistical analysis was completed using GraphPad Prism (v.10.6.1).

### Antibodies

| **Antigen** | **Host Species** | **Manufacturer** | **Catalog number** | **Application** | **Conjugation** |
| --- | --- | --- | --- | --- | --- |
| ATP5PB | Mouse | Abcam | Ab117991 | WB, Co-IP | - |
| ATP5MC1 | Rabbit | Abcam | Ab210732 | FC, IF | Alexa Flour 555 |
| ATP5F1B | Mouse | Abcam | Ab197649 | FC, IF | Alexa Flour 647 |
| ACTB | Mouse | Invitrogen | MA1-140 | WB | - |
| Rapsyn | Rabbit | Abcam | Ab156002 | WB, Co-IP | - |
| SLC25A1 | Rabbit | Proteintech | 15235-1-AP | WB | - |
| N-Cadherin | Rabbit | Abcam | Ab18203 | WB | - |
| HIF-1alpha | Rabbit | Cell Signaling | 14179 | WB | - |

## Results

### Cell surface Complex V expression

The presence of ectopic ATP synthase (ecto-ATP synthase) on the cell surface has been reported across multiple malignant cell types^26,33,38,39^, and it has been proposed that other complexes of the mitochondrial respiratory chain (MRC) may also localize to the plasma membrane^40,41^. To examine this, we probed non-permeabilized live primary retinoblastoma (Rb) cells with an antibody against the c-ring subunit of the F_O_ domain (ATP5MC1) of Complex V and counterstained the plasma membrane with CellMask Deep Red to assess co-localization by confocal microscopy (Fig. 1A). In RB006 cells, 25.7% of ATP synthase signal colocalized with the plasma membrane under normoxic conditions (21% O_2_), increasing to 64.6% after 12 hours of hypoxia (1% O_2_) (*p<0*.*0001*) (Fig. 1B). Similar plasma membrane localization and hypoxia-dependent increases were observed in RB036 primary cells expressing a ATP5F1B-EGFP fusion (Fig. S1A). Surface localization of complex V further increased in a time-dependent manner under 1% O_2_ (obligate glycolytic conditions) (Fig. S1B-D), while total cellular ATP synthase levels remained unchanged, consistent with a model of redistribution of pre-assembled complexes rather than increased de novo synthesis (Fig. 1C). In contrast, control HEK293 cells exhibited negligible surface ATP synthase signal and no response in hypoxic conditions (Fig.S2A,B).

**Fig. 1.**
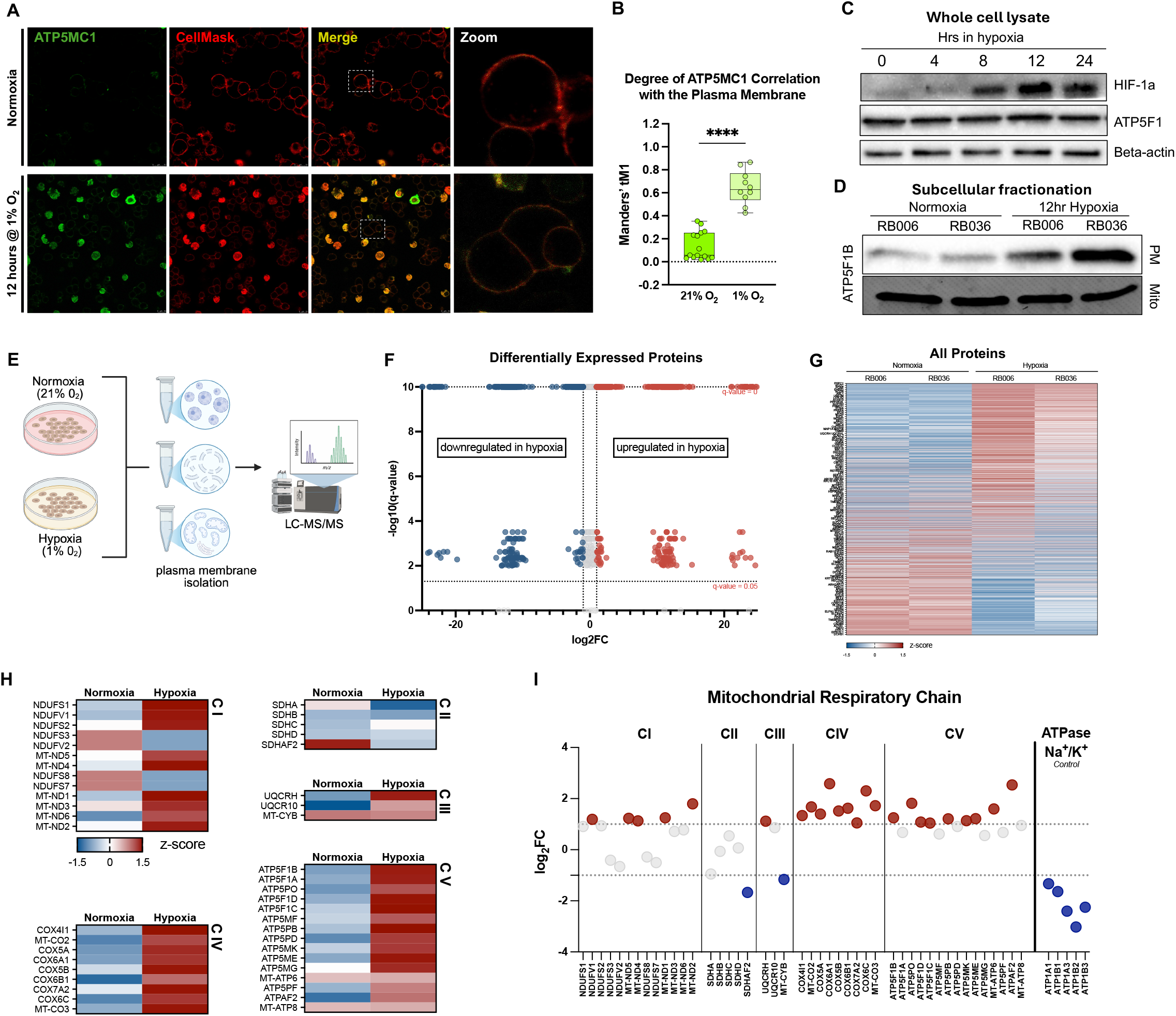
Oxidative phosphorylation machinery accumulates at the plasma membrane under glycolytic stress. **a**, Representative live-cell confocal images of non-permeabilized RB006 retinoblastoma cells stained for the ATP synthase F_O_ c-ring subunit ATP5MC1 and counterstained with CellMask to delineate the plasma membrane under normoxia (21% O2) or after 12 h hypoxia (1% O2). Insets show magnified regions. Scale bar, 10µm. **b**, Quantification of ATP5MC1 colocalization with the plasma membrane by Manders’ tM1 coefficient. Each point represents one image (*n = 10*). Representative of three biologically independent experiments. **c**, Immunoblot analysis of whole-cell lysates from RB006 and RB036 cells cultured under hypoxia for the indicated durations, probed for ATP5F1 and HIF-1α. Beta-actin served as loading control. Representative of three biologically independent experiments. **d**, Immunoblot analysis of isolated mitochondrial and plasma membrane fractions from RB006 and RB036 cells cultured under normoxia or after 12 h hypoxia, probed for ATP5F1B. Fraction purity controls are shown in Supplementary Fig. 3. Representative of three biologically independent experiments. **e**, Experimental workflow for plasma membrane isolation and LC–MS/MS proteomic profiling. **f**, Volcano plot showing differentially abundant proteins identified in isolated plasma membrane fractions under hypoxia relative to normoxia. Differential abundance thresholds were defined as log2 fold change ±1 and false discovery rate (FDR) <0.05. **g**, Heatmap of differentially abundant proteins identified in isolated plasma membrane fractions under normoxia and hypoxia. **h**, Heatmap of mitochondrial respiratory chain proteins identified in isolated plasma membrane fractions under normoxia and hypoxia. **i**, Differential abundance analysis of mitochondrial respiratory chain proteins identified in isolated plasma membrane fractions. *\*\*\*\* P < 0*.*0001* **(b)**. Statistics: two-tailed Welch’s t test **(b)**. Normalized z-scores **(g, h)**.

Biochemical fractionation followed by immunoblotting confirmed the presence of ATP synthase in purified plasma membrane fractions of Rb cells, with marked enrichment under hypoxia (Fig. 1D, Fig. S3), whereas HEK293 cells lacked detectable plasma membrane ATP synthase (Fig. S2C). Proteomic analysis of plasma membrane fractions by LC-MS/MS revealed a divergent protein composition profile between normoxic and hypoxic cells, with 508 (18.24%) proteins upregulated (log2FC > 1) and 435 (15.61%) proteins downregulated (log2FC < −1) in hypoxia (Fig. 1E-G, Fig. S4A,B). Hypoxic plasma membrane preparations were enriched for proteins of canonical mitochondrial localization, with inner mitochondrial membrane proteins comprising the largest fraction (31.6%)(Fig. S4C-H). Components of all MRC complexes (I-V) were detected within the plasma membrane and increased under hypoxia, with the most prominent enrichment noted in Complexes I, III, and IV, which all function as proton extruders and would contribute to the generation of an extracellular proton gradient at the plasma membrane (Fig. 1H, I, Fig. S4I). These data demonstrate glycolytic-dependent enrichment of ATP synthase and MRC components at the plasma membrane of Rb cells.

### Ecto-ATP synthase generates intracellular ATP

To determine whether ecto-ATP synthases are oriented to support intra- or extracellular ATP production, we assessed the accessibility of the F_O_ and F_1_ domains to the extracellular space using a nonlytic HiBiT-based detection system (Fig. 2A). RB006, RB036, and control mesenchymal stem cells (MSCs) were engineered to express either ATPME-HiBiT or ATP5F1B-HiBiT, effectively tagging either the e-subunit of the F_O_ domain or the beta subunit of the F_1_ domain, respectively. Use of a nonlytic buffer in conjunction with the non-cell permeable Large BiT (LgBiT) subunit, allows for the sensitive detection of the HiBiT-tagged protein that is accessible to the extracellular space. Hypoxia induced a 2.8- and 2.5-fold increase in extracellular F_O_ signal in RB006 and RB036, respectively (*p<0*.*0001*)(Fig. 2B), compared to more modest increases in F_1_ exposure (FC = 1.3 and 1.4, respectively; *p<0*.*001* and *p<0*.*01*, respectively) (Fig. 2C). When normalized to total cellular ATP synthase, quantified using a lytic detection reagent to measure intracellular HiBiT-tagged proteins, hypoxia increased surface localization of the F_O_ domain from 20% to 68%, compared to an increase in F_1_ from 8% to 20% (*p<0*.*0001*) (Fig. 2D). Control MSCs showed negligible surface expression of either domain, and no response to hypoxia. This preferential extracellular exposure of the F_O_ domain under hypoxia is consistent with an orientation that supports intracellular ATP production.

**Fig. 2.**
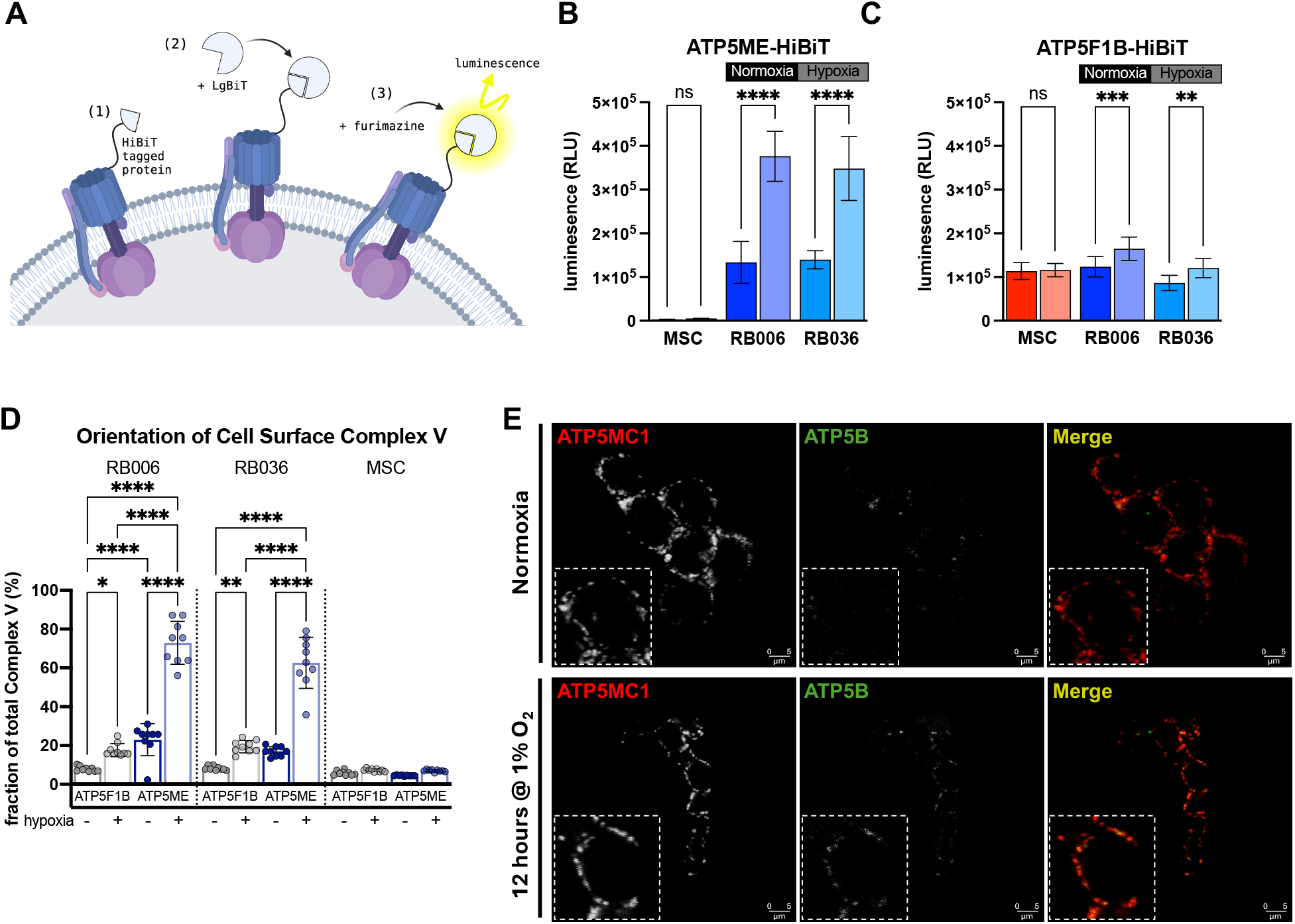
Cell surface ATP synthase adopts an F_O_-outward orientation. **a**, Schematic of the non-lytic HiBiT/LgBiT assay used to assess extracellular accessibility of ATP synthase domains. ATP5ME-HiBiT and ATP5F1B-HiBiT constructs were used to label the F_O_ and F_1_ domains, respectively. **b**, Extracellular luminescence signal from ATP5ME-HiBiT-expressing mesenchymal stem cells (MSCs), RB006 and RB036 cells cultured under normoxia or after 12 h hypoxia. Data are from three biologically independent experiments. **c**, Extracellular luminescence signal from ATP5F1B-HiBiT-expressing MSCs, RB006 and RB036 cells cultured under normoxia or after 12 h hypoxia. Data are from three biologically independent experiments. **d**, Quantification of extracellularly accessible ATP synthase normalized to total cellular ATP synthase abundance, calculated as non-lytic luminescence relative to total lytic luminescence. Data are from three biologically independent experiments. **e**, Representative live-cell confocal images of non-permeabilized RB036 cells stained with antibodies against the ATP synthase F_O_ domain (ATP5MC1) and F_1_ domain (ATP5B) under normoxia or after 12 h hypoxia. Scale bar, 5 µm. * *P <0*.*05, ** P < 0*.*01, *** P <0*.*001, **** P < 0*.*0001* normoxia versus hypoxia **(b, c)** or all conditions within a cell type **(d)**. Statistics: two-tailed Welch’s t test **(b, c);** two-way ANOVA with Šídák multiple comparisons test **(d)**. Error bars: mean ± s.d.

Orientation was further assessed using domain-specific antibodies in non-permeabilized cells (Fig. 2E). Both domains were detectable on the cell surface; however, the F_O_ domain was the predominantly extracellularly exposed component, whereas F_1_ staining appeared sparser (Fig. 2E). Flow cytometry confirmed surface expression of both domains in Rb cells (Fig. S5A), with a preferential increase in extracellular F_O_ domain exposure under hypoxia. The proportion of cells with extracellular F_O_ increased from 41.9% to 51.8% in RB006 (*p<0*.*01*) and from 26.1% to 48.9% in RB036 (*p<0*.*0001*), whereas extracellular F_1_ decreased in both lines under hypoxia (Fig. S5B). Consequently, the F_O_:F_1_ orientation ratio shifted from 0.69 under normoxia to approximately 1.4 under hypoxia (Fig. S5C), suggesting preferential reorientation toward an F_O_-outward configuration.

### Respiratory complexes embed via fusion

Our proteomic analysis identified several mitochondrial proteins within the plasma membrane fraction (Fig. S4), suggesting a mechanism involving mitochondrial trafficking to, and fusion with, the plasma membrane. Live-cell imaging of Rb cells expressing mitochondria-targeted mCherry revealed punctate mitochondrial localization at the plasma membrane, counterstained with CellMask Deep Red, which was increased under hypoxia (mean difference of ~8%; *p<0*.*001*) (Fig. 3A,B). Conversely, co-expression of ATP5F1B-EGFP and mitochondria-targeted mCherry in HEK293 showed no substantial localization or fusion with the plasma membrane (Fig. S6A,B, Fig, S7). Consistent with these observations, transmission electron microscopy (TEM) revealed mitochondria in close apposition to the plasma membrane in cancer cells, forming direct contact points, membrane bridges, and structures consistent with fusion intermediates (Fig. 3C, Fig. S8A). Intact mitochondria and mitochondrial remnants also appeared within budding blebs, as also observed by live-cell confocal microscopy (Fig. S8B,C), supporting the premise of active mitochondrial recruitment to the cell periphery and providing an intermediate through which mitochondrial membranes may dock, remodel, or partially fuse with the plasma membrane. These findings support mitochondrial trafficking and fusion as a mechanism for incorporation of ATP synthase complexes and associated components into the plasma membrane.

**Fig. 3.**
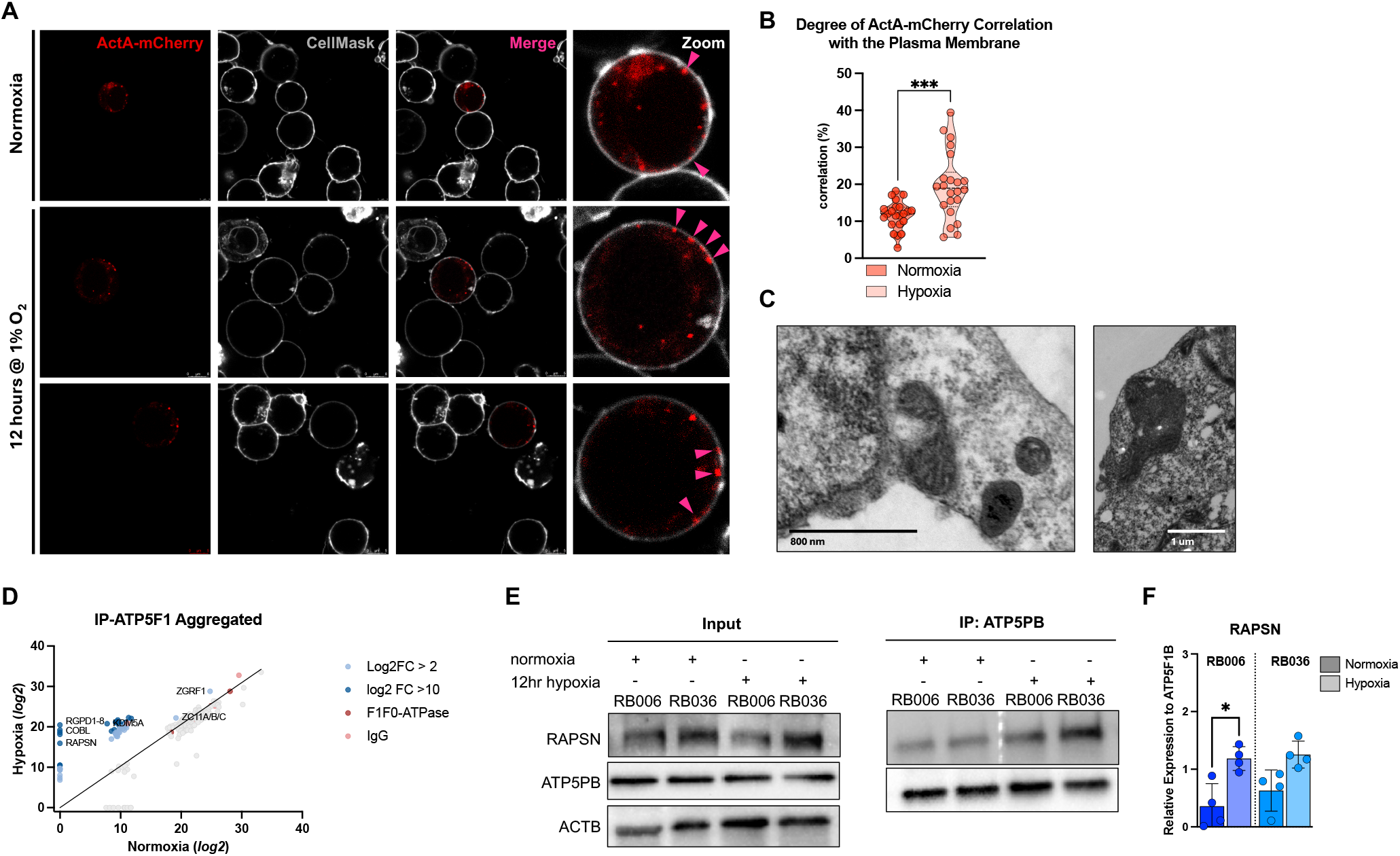
Mitochondrial trafficking mediates plasma membrane incorporation of oxidative phosphorylation machinery. **a**, Representative live-cell confocal images of RB006 cells expressing mitochondria-targeted ActA-mCherry and counterstained with CellMask to delineate the plasma membrane under normoxia (21% O2) or after 12 h hypoxia (1% O2). Insets show magnified regions. Scale bar, 5 µm. **b**, Quantification of mitochondrial colocalization with the plasma membrane. Each point represents an individual mitochondrion. Data are pooled from ten images acquired from three biologically independent experiments. **c**, Representative transmission electron micrographs of retinoblastoma cells. Representative of three biologically independent experiments. Scale bar, X µm. **d**, Scatter plots showing proteins identified by ATP5PB co-immunoprecipitation followed by mass spectrometry in RB006 and RB036 cells under normoxia or after 12 h hypoxia. **e**, Immunoblot validation of ATP5PB co-immunoprecipitation with RAPSN in RB006 and RB036 cells under normoxia or after 12 h hypoxia. Representative of three biologically independent experiments. **f**, Quantification of ATP5PB–RAPSN co-immunoprecipitation signal from three biologically independent experiments. * *P <0*.*05, *** P <0*.*001* normoxia versus hypoxia **(b, f)**. Statistics: two-tailed Welch’s t test **(b, f)**.

To identify potential mechanisms of ecto-ATP synthase anchoring within the plasma membrane, we performed co-immunoprecipitation of the ATP synthase peripheral stalk-membrane subunit b (ATP5PB) followed by mass spectrometry. Among 197 unique interacting proteins, rapsyn was identified as a hypoxia-specific binding partner in both RB006 and RB036 (Fig. 3D). This interaction was validated by immunoblot (Fig. 3E, F). Rapsyn canonically anchors acetylcholine receptors (AChR) at the plasma membrane, yet no AChRs were detected in the plasma membrane of our cells, suggesting a previously unrecognized role for rapsyn in stabilizing ecto-ATP synthase complexes in the plasma membrane.

### Ecto-ATP synthases respond to extracellular acidity

With ATP synthase localized to the cell surface and predominantly oriented with the F_O_ domain facing the extracellular space, we investigated whether extracellular acidification influences ATP production. Prior studies have demonstrated that ecto-ATP synthase retains enzymatic activity at the cell surface^26–33,42–46^, although these studies primarily report the production of extracellular ATP. In contrast, we posit that cell surface ATP synthase can drive intracellular ATP, fueled by the proton gradient established by lactate export and subsequent extracellular acidity.

Cells were cultured under progressively acidic conditions and both intracellular and extracellular ATP levels were quantified. In Rb cells, intracellular ATP production increased in both a pH- and time-dependent manner, rising with decreasing extracellular pH (Fig. 4A, B). After 60 minutes, ATP levels remained unchanged under physiological pH (fold change = 1.0), whereas cells cultured at pH 6 and pH 5 exhibited a 1.4- and 1.8-fold increase in intracellular ATP, respectively (*p<0*.*0001*) (Fig. 4C). RB036 demonstrated the most pronounced response, with a near-linear increase in ATP production as pH decreased beginning at 15 minutes and persisting throughout the assay, whereas RB006 showed a more modest but still consistent increase. Under hypoxia, ATP production was further augmented by the acidic microenvironment, increasing by 1.6-fold at pH 6 and 2.3-fold at pH 5, while intracellular ATP levels under physiological pH conditions declined by approximately 20% over time (Fig. 4D-F). In contrast, MSCs showed no consistent increase in intracellular ATP production under either normoxic or hypoxic conditions (Fig. S9A-F).

**Fig. 4.**
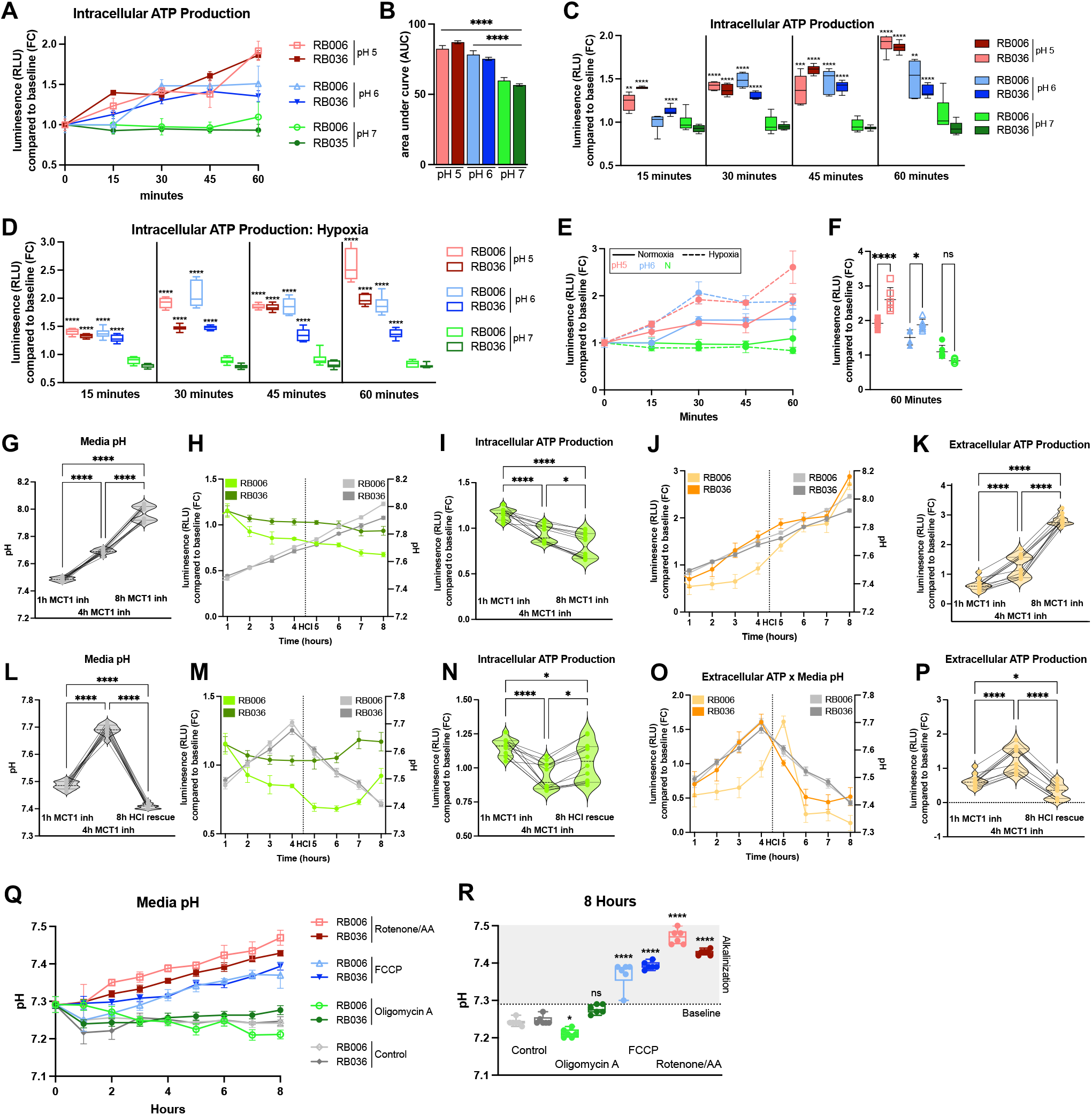
Extracellular proton gradients drive ATP production through cell surface ATP synthase. **a**, Time-course quantification of intracellular ATP in RB006 and RB036 cells cultured in 5 mM HEPES-buffered media at pH 7.4, pH 6 or pH 5 over 60 min, normalized to baseline (t = 0). Data are from three biologically independent experiments. **b**, Area under the curve (AUC) analysis of intracellular ATP under the indicated extracellular pH conditions. **c**, Comparison of intracellular ATP at 15-, 30-, 45-, and 60 min across extracellular pH conditions. Data are from three biologically independent experiments. **d**, Comparison of intracellular ATP at 15-, 30-, 45-, and 60 min across extracellular pH conditions under hypoxia (1% O_2_). Data are normalized to baseline (t = 0). Data are from three biologically independent experiments. **e**, Time-course comparison of intracellular ATP in RB006 and RB036 cells cultured in 5 mM HEPES-buffered media at pH 7.4, pH 6 or pH 5 over 60 min, normalized to baseline (t = 0). Cells were cultured in either normoxic (solid line) or hypoxic (dashed line) conditions. Data are from three biologically independent experiments. **f**, Comparison of intracellular ATP at 60 min across extracellular pH conditions under normoxia (filled shapes) and hypoxia (open shapes). Data are from three biologically independent experiments. **g**, Media pH in RB006 and RB036 cells treated with AZD3965 (20 nM) over 8 h. Data are from three biologically independent experiments. **h**, Time-course of intracellular ATP (left axis) and media pH (right axis) in RB006 and RB036 cells following AZD3965 treatment. Intracellular ATP production is normalized to baseline (t = 0). Data are from three biologically independent experiments. **i**, Quantification of intracellular ATP following AZD3965 treatment at indicated time points. Data are from three biologically independent experiments. **j, k**, Same as in **(h, i)** for quantification of extracellular ATP production. **l-p**, Same as in **(g-k)**; after 4 hours of AZD3965 treatment, media was re-acidified with HCl (5mM). Data are from three biologically independent experiments. **q**, Measurement of media pH over 8 h following treatment with rotenone/antimycin A (0.125 µM), oligomycin A (0.375 µM), or FCCP (0.125 µM). Data are from three biologically independent experiments. **r**, Endpoint quantification of extracellular media pH following treatment with rotenone/antimycin A, oligomycin A or FCCP. * *P <0*.*05, ** P < 0*.*01, **** P < 0*.*0001* versus normal media pH **(b-d)**; normoxia versus hypoxia **(f)**; comparison of each time point **(g, i, k, l, n, p)**; versus control **(r)**. Statistics: one-way ANOVA with Tukey’s multiple comparisons test **(b, d, f, g, i, k, l, n, p, r)**. Error bars: mean ± s.d. **(a, d, e, h, j, m, o, q)**; box and whisker plots: min-to-max **(c, d, r)**.

In contrast, extracellular ATP levels decreased with increasing extracellular acidity, with the most pronounced effect seen in RB036. Overall, extracellular ATP production was reduced by approximately 8% at pH 6 (*p < 0*.*0001*) and 32% at pH 5 (*p < 0*.*0001*), whereas cells cultured under physiologic pH had no change compared to baseline (Fig. S10A-C). Under hypoxia, this effect was more pronounced, with a 17% and 29% reduction at pH 5 and 6 (*p<0*.*0001*), respectively, and a nearly 10% increase at physiological conditions (*p=0*.*0002*) (Fig. S10D-F). MSCs again showed no change in extracellular ATP production in response to extracellular pH under any condition (Fig. S9G-L).

Lactate export via monocarboxylate transporters (MCTs) is a reported leading contributor to extracellular acidification in cancer cells, stemming from their high glycolytic rate. In retinoblastoma, MCT1 is the primary monocarboxylate transporter, and its expression has been correlated with prognosis.^47^ To assess the contribution of extramitochondrial ATP production by lactate export, cells were treated with the monocarboxylate transporter 1(MCT1) inhibitor AZD3965 (20nM), resulting in a progressive increase in media pH, consistent across all Rb cell lines (*p<0*.*0001*) (Fig. 4G). This alkalinization was accompanied by a concomitant reduction in intracellular ATP production, showing a 32.3% and 6.5% decline in RB006 (*p<0*.*0001*) and RB036 (*p=0*.*020*), respectively, alongside a nearly threefold increase in extracellular ATP production after 8 hours (*p<0*.*0001*)(Fig. 4H-K). Under hypoxic conditions, intracellular ATP production in response to MCT1 inhibition was significantly augmented, with a 44% average decrease across RB006 and RB036 (*p<0*.*0001*), whereas extracellular ATP production did not change significantly compared to normal oxygen conditions (Fig. S11A-F). MSCs showed no significant changes in either intracellular or extracellular ATP levels (*p=0*.*974; p=0*.*986*, respectively) (Fig. S12A-E).

Re-acidification of the media with HCl (5 mM) following 4 h of AZD3965 treatment rescued intracellular ATP production from 7% below baseline to 5% above baseline in Rb cells (*p=0*.*019*), while also suppressing extracellular ATP levels from 26% above baseline to 67% below baseline (*p<0*.*0001*) (Fig. 4L-P). The same trend was observed under hypoxia, with intracellular ATP production being significantly augmented with re-acidification, showing a 26% increase (*p<0*.*0001*), compared to the 12% observed under normoxia (*p=0*.*0001*) (Fig. S11G-L). MSCs remained unresponsive to MCT1 inhibition, despite concordant changes in media pH (Fig. S12F-J). Together, these results indicate that extracellular pH and lactate transport are critical in modulating intra- and extracellular ATP production in an inverse manner.

Provided that proton effluxing mitochondrial respiratory chain complexes were also found to be embedded within the plasma membrane (Fig. 1H), we examined their potential contribution to extracellular acidification with ultra-low-dose inhibition of Complexes I (Rotenone), III (Antimycin A), V (Oligomycin A), and the protonophore carbonyl cyanide-p-trifluoromethoxyphenylhydrazone (FCCP) (Fig. S13). Inhibition of Complexes I and III, known proton transporters, with combination rotenone/Antimycin A resulted in the largest increase in extracellular pH, with an average rise of 0.205 units over 8 hours (*p < 0*.0001), followed by a 0.137 unit rise with non-specific disruption of the proton gradient with FCCP (*p<0*.*0001*) (Fig. 4Q, R). These results confirm the functionality of embedded mitochondrial respiratory chain complexes and their limited contribution to extracellular acidification. Oligomycin A, on the other hand, resulted in no change in pH compared to control, further supporting our findings that ecto-ATP synthases are oriented with the F_O_ domain in the extracellular space (Fig. 2), functioning to transport protons towards the intracellular space.

### F_O_ domain is a targetable vulnerability

To assess the therapeutic targetability of the extracellular F_O_ domain of ecto-ATP synthase, we evaluated the effects of the F_O_ proton channel inhibitor dibutyltin-3-hydroxyflavone chloride [Bu_2_Sn(of)Cl]^48^ in proof-of-concept dose-dependent viability assays. Rb and mesenchymal stem cells (MSCs) were treated for 72 hours and analyzed using MTT and CellTiter-Glo 2.0. In the MTT assay, Bu_2_Sn(of)Cl reduced viability of RB006 and RB036 in a dose-dependent manner, with a half maximal inhibitory concentration (IC_50_) of 610 nM and 447 nM, respectively (Fig. 5A, B). In contrast, MSCs maintained greater than 50% viability at all concentrations tested, with a calculated IC_50_ of 11.5 µM (Fig. 5C), and significant differences in viability between Rb and MSCs at all doses (Fig. 5D). CellTiter-Glo 2.0 assays showed a pronounced therapeutic index in cancer cells, with an average IC_50_ of 0.147 µM and 0.289 µM in RB006 and RB036, respectively, compared to 76.83µM in MSCs (*p<0*.*0001*) (Fig. 5E-H). Bu_2_Sn(of)Cl also induced dose-dependent cell death in a human neuroblastoma cell line (SH-SY5Y), which has been reported to expresses ecto-ATP synthase^49,50^ (Fig. S14A, B). To determine whether hypoxia alters inhibitor sensitivity, cells were treated under hypoxic conditions. Hypoxia produced a slight but non-significant rightward shift of the viability curves across all cell lines (Fig. S14C-F), consistent with our observations of a higher abundance of ecto-ATP in glycolytic obligate cancer cells. These results show selective sensitivity of cancer cells to F_O_ domain inhibition.

**Fig. 5.**
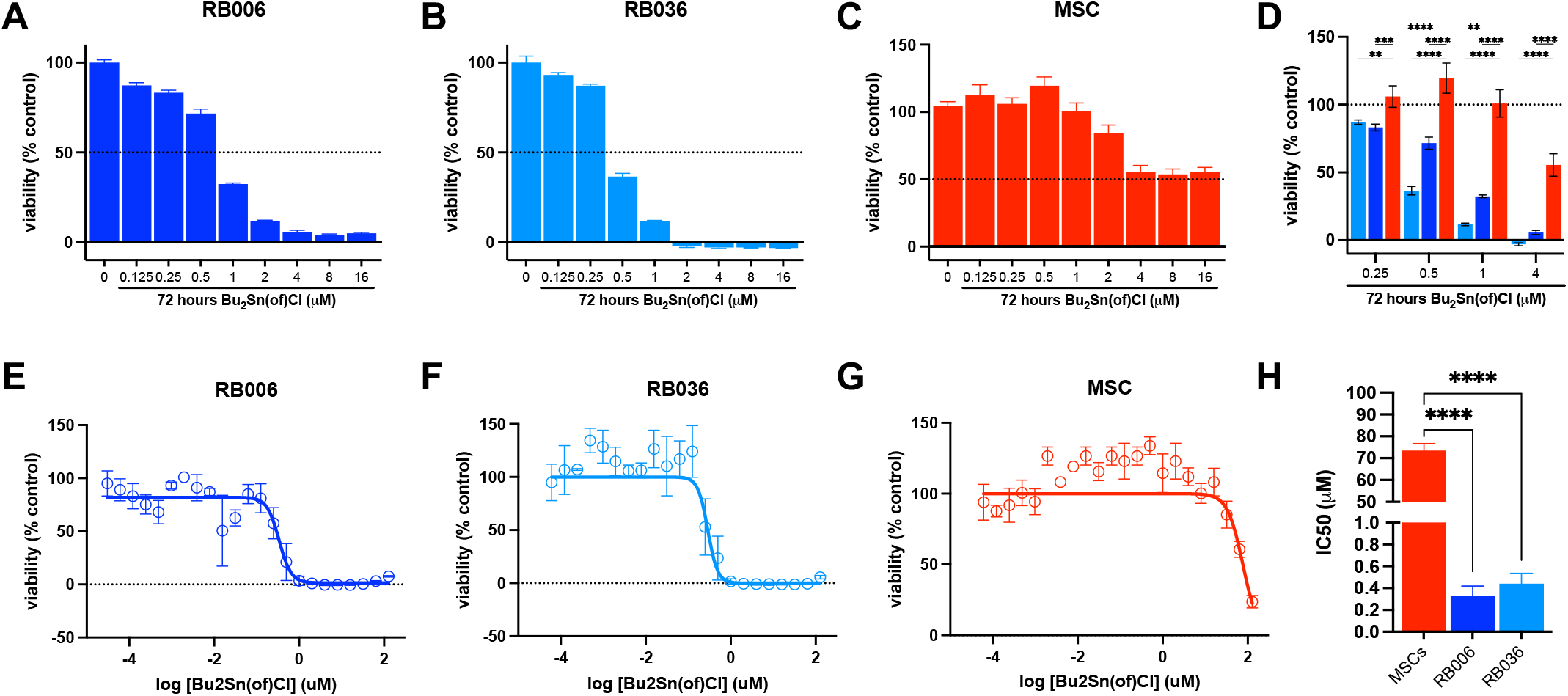
The extracellular FO domain of ATP synthase is a targetable metabolic vulnerability. **a-c**, Dose–response analysis of RB006, RB036, and MSC cell viability, respectively, following 72 h treatment with dibutyltin-3-hydroxyflavone chloride (Bu2Sn(of)Cl), quantified by MTT assay. Data are from three biologically independent experiments. **d**, Quantification of cell viability across the indicated Bu2Sn(of)Cl concentrations in RB006, RB036 and MSCs measured by MTT assay. *(n = 3*) **e-g**, Dose–response analysis of RB006, RB036, and MSC viability, respectively, following 72 h Bu2Sn(of)Cl treatment, quantified by CellTiter-Glo 2.0 assay. Data are from three biologically independent experiments. Nonlinear regression was performed using a four-parameter variable-slope model. **h**, Quantification of Bu2Sn(of)Cl IC_50_ in RB006, RB036 and mesenchymal stem cells measured by CellTiter-Glo 2.0 assay (*n = 3*). *\*\* P < 0*.*01, *** P <0*.*001, **** P < 0*.*0001* cell viability at each dose **(d)** and comparing IC_50_ **(h)**. Statistics: two-way ANOVA with Šídák multiple comparisons test **(d)**; one-way ANOVA with Tukey’s multiple comparisons test **(h)**. Error bars: mean ± s.d..

## Discussion

The Warburg effect has long been recognized as a defining feature of cancer metabolism, yet its energetic teleology remains unresolved. Although aerobic glycolysis has been linked to biosynthesis^1,8,9^, redox balance^14,15^, hypoxic adaptation^16^, and microenvironmental remodeling^24,25^, these frameworks do not fully reconcile a central paradox: why cancer cells preferentially utilize a metabolically inefficient pathway that yields substantially less ATP per molecule of glucose than oxidative phosphorylation. The conservation of this phenotype across tumor types suggests that aerobic glycolysis confers a previously unrecognized energetic advantage.

Here, we redefine aerobic glycolysis as a bioenergetic strategy that directly contributes to ATP production through spatial reorganization of the oxidative phosphorylation machinery. Specifically, we identify a mechanism linking a defining feature of the Warburg phenotype - lactate-driven extracellular acidification - to ATP production via a plasma membrane-localized ATP synthase (Complex V). We show that translocated ATP synthase complexes embedded in the plasma membrane can exploit the extracellular proton gradient generated by lactate export to drive intracellular ATP synthesis outside of mitochondria. These findings extend prior observations of cell-surface ATP synthase^26,33,38,39^ and respiratory chain components^39–41^ by demonstrating that this system is catalytically active and contributes directly to intracellular ATP pools.

This conceptual framework is supported by a structural model in which mitochondrial components, including ATP synthase and proton-translocating respiratory complexes, are trafficked to and integrated within the plasma membrane through mitochondrial fusion-associated processes. This redistribution establishes a structural basis for an extramitochondrial, proton-coupled ATP-generating system at the cell surface. In this configuration, extracellular acidification establishes a proton gradient across the plasma membrane analogous to the mitochondrial proton motive force, while the presence of respiratory chain components may further reinforce this gradient. Notably, this relocalization is selectively enriched under hypoxic conditions in cancer cells but not in non-malignant proliferative cells, suggesting that it represents a cancer-specific metabolic plasticity associated with glycolytic dependence.

Within this context, plasma membrane ATP synthase can be understood as a bidirectional, proton gradient-dependent enzyme whose catalytic direction is governed by proton gradient polarity. Under conditions of extracellular acidification characteristic of the tumor microenvironment, proton influx through the F_O_ domain favors intracellular ATP synthesis, whereas reversal of the gradient can promote extracellular ATP release. This framework reconciles prior reports of extracellular ATP production^27,41,51–54^ with our observation that intracellular ATP generation predominates under physiologically relevant tumor conditions.

The biochemical constraints of the tumor microenvironment further support this interpretation. Elevated extracellular proton concentration, together with a relatively lower resting plasma membrane potential compared to the inner mitochondrial membrane^55,56^, would render sustained extracellular ATP synthesis thermodynamically unfavorable. At the same time, persistent extracellular ATP accumulation is difficult to reconcile with the immunosuppressive nature of tumors given the well-established role of extracellular ATP as an immunogenic signal^57–60^. These considerations converge on a model in which intracellular ATP production represents the dominant physiological function of cell surface ATP synthase in cancer.

This framework provides a potential resolution to the longstanding energetic paradox of the Warburg effect. Rather than representing a metabolically inefficient state, aerobic glycolysis may function to generate an extracellular proton gradient that can be harnessed for ATP production. In this context, glycolysis, lactate export, extracellular acidification, and ATP synthesis operate as components of an integrated bioenergetic architecture in cancer, rather than independent metabolic processes. This perspective offers a mechanistic explanation for the consistent selection of high glycolytic flux in tumors despite its apparent inefficiency in isolation.

The ability to spatially reorganize bioenergetic machinery in this manner points to a broader capacity for structural and metabolic plasticity in cancer cells. Redistribution of mitochondrial components to the plasma membrane enables tumor cells to compensate for hypoxic, nutrient-limited, and acidic environments by exploiting extracellular conditions to sustain energy production, a property not observed in non-malignant proliferative cells.

These findings also suggest a potential therapeutic vulnerability. The extracellular F_O_ domain of ATP synthase, positioned to mediate proton flux across the plasma membrane, represents a functionally critical component of this energetic architecture. While prior efforts have focused on the F_1_ domain^29,32–34,38,61,62^, targeting the F_O_ proton-conducting domain may provide a more direct means of disrupting this bioenergetic axis. The preferential cytotoxicity in cancer cells is consistent with their reliance on this mechanism, although further optimization will be required to improve specificity and limit toxicity of F_O_-targeting compounds.

In summary, we define a previously unrecognized bioenergetic axis in cancer, in which lactate-driven extracellular acidification is coupled to ATP production through plasma membrane-localized ATP synthase. This mechanism provides a functional resolution to the longstanding paradox of the apparent energetic inefficiency of the Warburg effect and suggests that cancer metabolism involves not only metabolic reprogramming but also spatial reorganization of bioenergetic machinery. Determining the extent to which this pathway operates across tumor types and contributes to tumor progression, therapeutic resistance, or metabolic adaptability will be an important direction for future investigation.

## Acknowledgements

We thank the donors and their families for their extensive contribution to science. We acknowledge the support of the Miami Project to Cure Paralysis Transmission Electron Microscopy Core Facility (Vania Almeida) and the Sylvester Comprehensive Cancer Center Flow Cytometry Shared Resource. This work has been supported in part by the Proteomics and Metabolomics Core Facility at the H. Lee Moffitt Cancer Center & Research Institute; an NCI designated Comprehensive Cancer Center (P30-CA076292).

