## Supplemental Figures for "Surface Complex V couples proton gradient across the plasma membrane to ATP production in cancer"

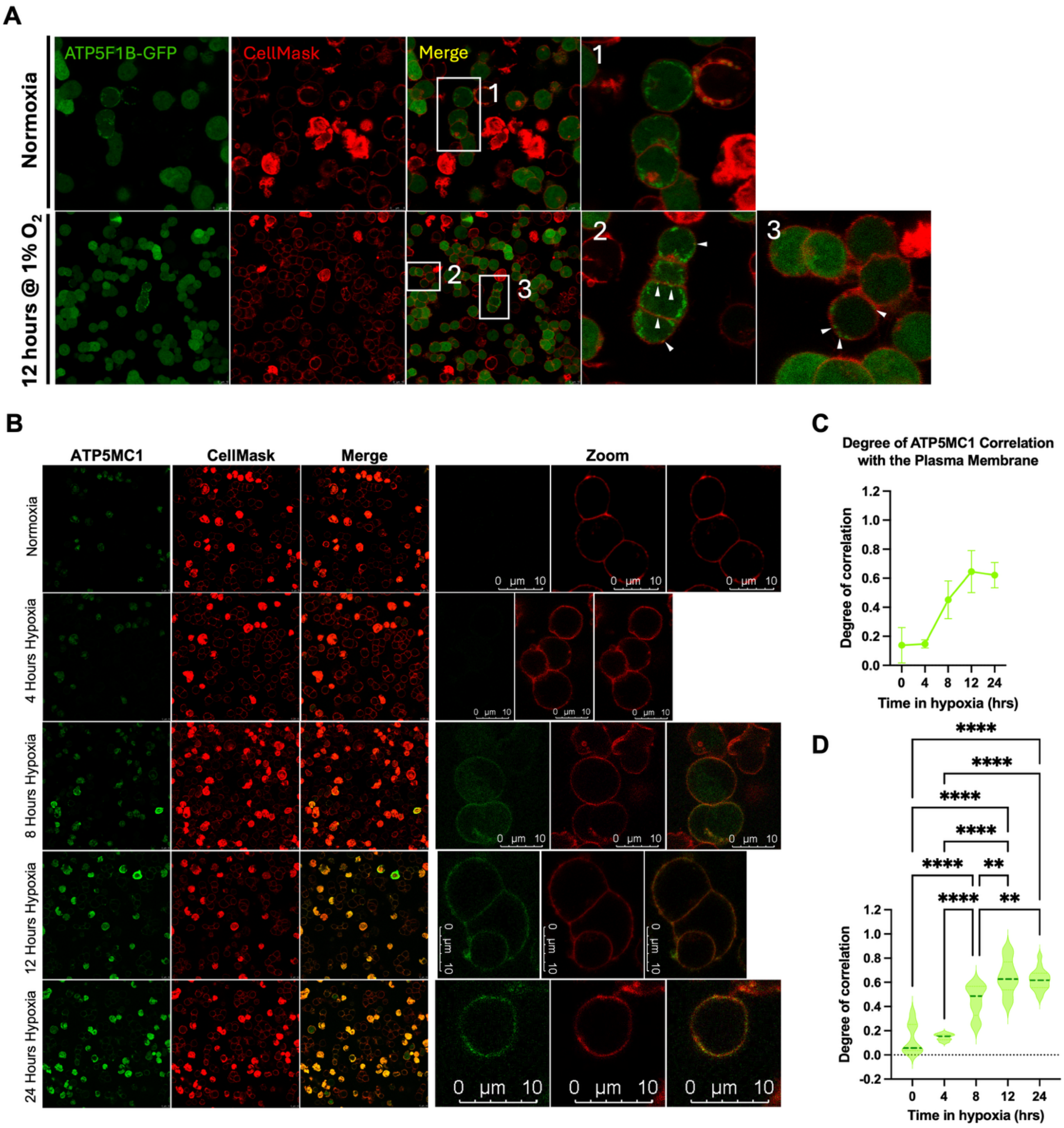


**Supplementary Fig. 1 | Hypoxia promotes plasma membrane localization of ATP synthase.** **a**, Representative live-cell confocal images of RB036 cells expressing ATP5F1B-EGFP and counterstained with CellMask under normoxia (21% $O_2$) or after 12 h hypoxia (1% $O_2$). Insets show magnified regions. Scale bar, 10 µm. **b**, Representative live-cell confocal images of non-permeabilized RB006 cells stained for ATP5MC1 and counterstained with CellMask following 0, 4, 8, 12 or 24 h hypoxia. Insets show magnified regions. Scale bar, 10 µm. **c**, Time-course quantification of ATP5MC1 colocalization with the plasma membrane by Manders' tM1 coefficient across hypoxic exposure durations. **d**, Quantification of ATP5MC1 colocalization with the plasma membrane at indicated hypoxic time points. Each point represents one image. Data are pooled from ten images acquired across three biologically independent experiments. ** $P < 0.01$, **** $P < 0.0001$. Statistics: one-way ANOVA with Tukey's multiple comparisons test **(d)**. Error bars: mean ± s.d.

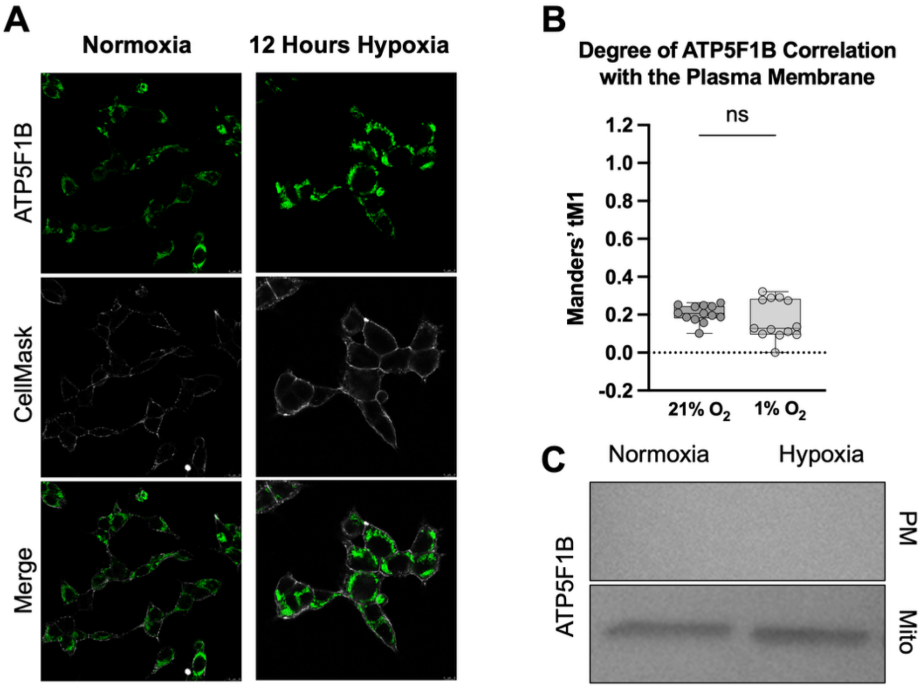


**Supplementary Fig. 2 | ATP synthase plasma membrane localization is not observed in HEK293 cells. a,** Representative live-cell confocal images of HEK293 cells expressing ATP5F1B-EGFP and counterstained with CellMask under normoxia or after 12 h hypoxia. Scale bar, 10 µm. **b,** Quantification of ATP5F1B-GFP colocalization with the plasma membrane by Manders' tM1 coefficient. Each point represents one image (n = 13). Representative of three biologically independent experiments. **c,** Immunoblot analysis of isolated mitochondrial and plasma membrane fractions from HEK293 cells cultured under normoxia or after 12 h hypoxia, probed for ATP5F1B. Representative of three biologically independent experiments. ns, not significant. Statistics: two-tailed Welch's *t*-test (c). Error bars: min-to-max.

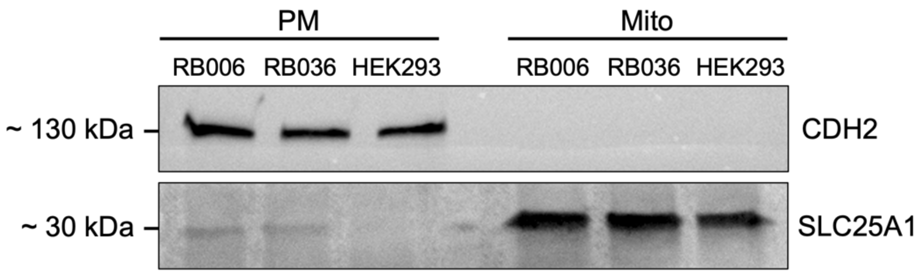


**Supplementary Fig. 3 | Validation of plasma membrane fraction purity.** Immunoblot analysis of isolated plasma membrane and mitochondrial fractions from RB006, RB036 and HEK293 cells probed for the plasma membrane marker CDH2 and mitochondrial marker SLC25A1. Representative of three biologically independent experiments.

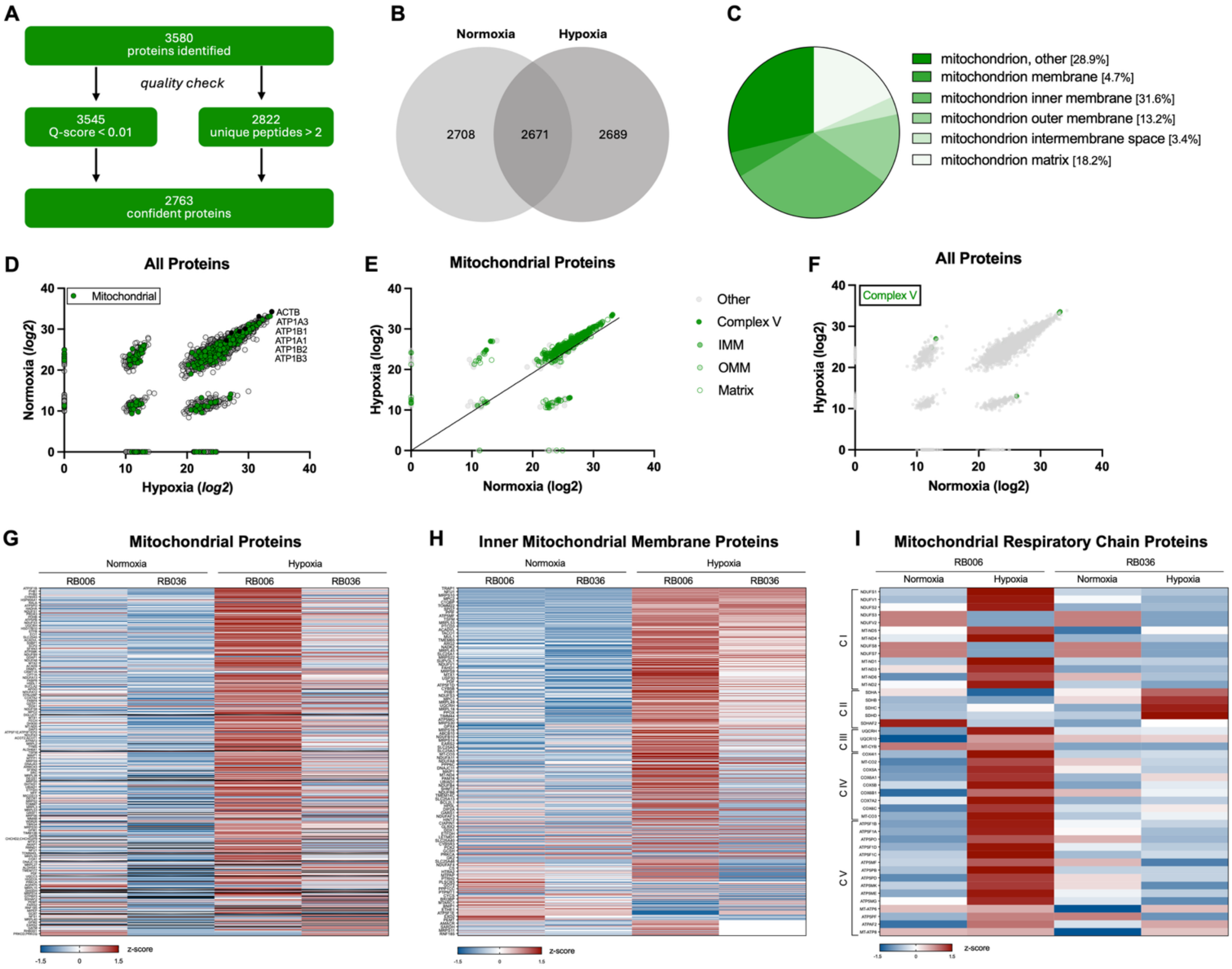


**Supplementary Fig. 4 | Proteomic characterization of isolated plasma membrane fractions.** **a,** LC–MS/MS proteomic workflow and quality control metrics for isolated plasma membrane fractions. **b,** Quantification of mitochondrial proteins identified in isolated plasma membrane fractions under normoxia and hypoxia. **c,** Quantification of subcellular location of mitochondrial proteins identified in isolated plasma membrane fractions. **d–e,** Mitochondrial proteins identified in isolated plasma membrane fractions and their relative expression under normoxia and hypoxia. **f,** Complex V expression in the plasma membrane identified under normal and hypoxic conditions. **g-i,** Heatmap depicting differential abundance of mitochondrial proteins, inner mitochondrial membrane proteins, and mitochondrial respiratory chain proteins identified in isolated plasma membrane fractions. Normalized z-scores **(g-i)**.

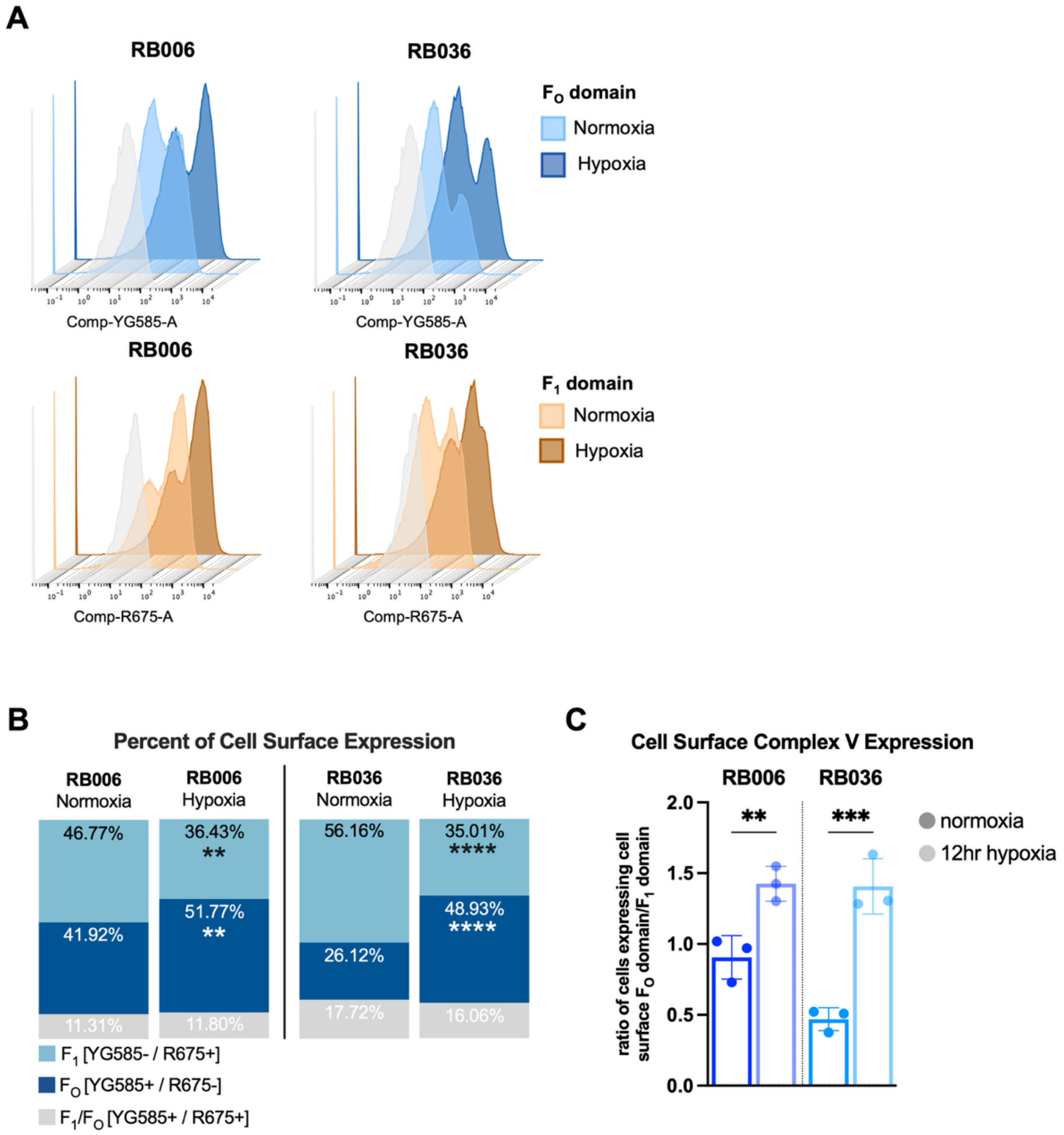


**Supplementary Fig. 5 | Cell surface accessibility of ATP synthase domains by flow cytometry. a,** Representative 3D flow cytometry histograms showing extracellular ATP synthase $F_O$ domain (ATP5MC1) and $F_1$ domain (ATP5B) in RB006 and RB036 cells cultured under normoxia (21% $O_2$) or after 12 h hypoxia (1% $O_2$). **b,** Quantification of the percentage of cells positive for cell surface ATP5MC1 and/or ATP5B under normoxia and hypoxia. Data are from three biologically independent experiments. **c,** Quantification of the extracellular $F_O$:$F_1$ domain ratio under normoxia and hypoxia. Data are from three biologically independent experiments. *** $P < 0.01$, *** $P < 0.001$, **** $P < 0.0001$* versus normoxia **(b, c)**. Statistics: two-tailed Welch's *t*-test **(b, c)**. Error bars: mean ± s.d.

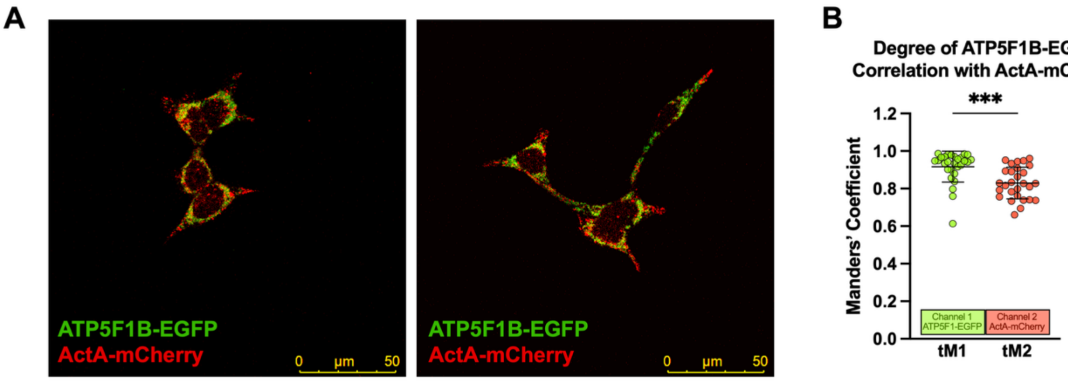


**Supplementary Fig. 6 | Validation of ActA-mCherry . a,** Representative confocal images of HEK293 cells expressing ATP5F1B-EGFP and mitochondria-targeted ActA-mCherry. **b,** Quantification of ATP5F1B-EGFP colocalization with mitochondria-targeted ActA-mCherry by Manders' coefficients (tM1 and tM2). Data are from three biologically independent experiments. *** $P < 0.001$. Statistics: two-tailed Welch's ***t***-test (b). Error bars: mean ± s.d.

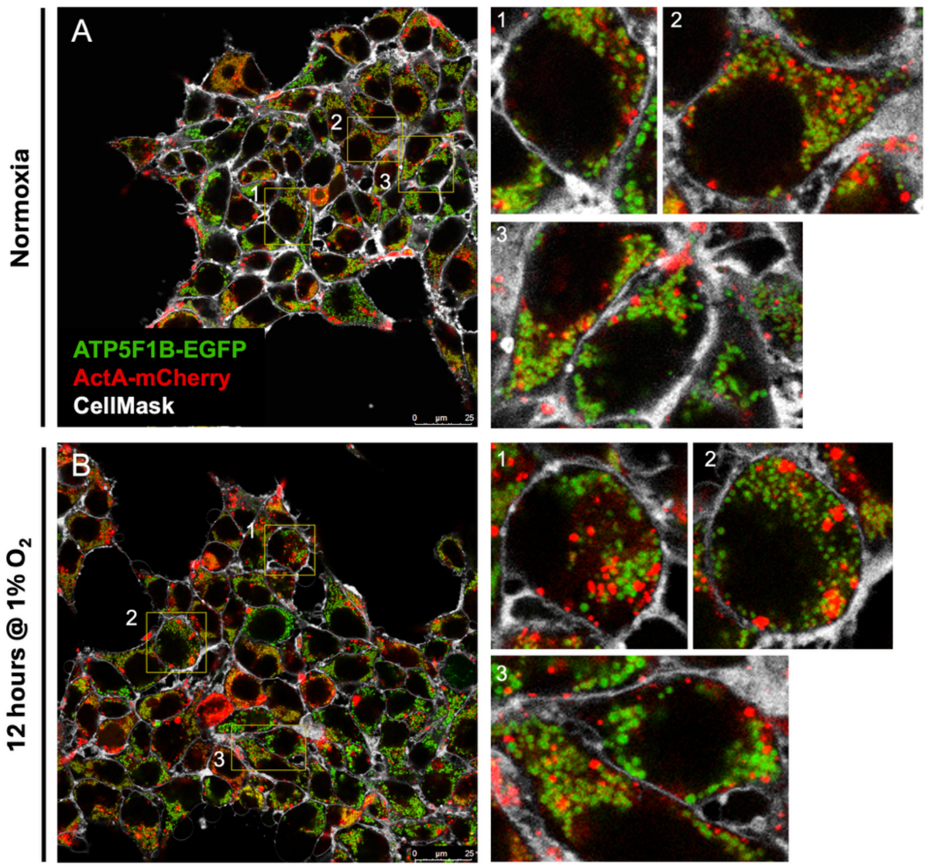


**Supplementary Fig. 7 | ATP synthase and mitochondrial localization in HEK293 cells.**
**a,** Representative live-cell confocal images of HEK293 cells expressing ATP5F1B-EGFP and mitochondria-targeted ActA-mCherry under normoxia or after 12 h hypoxia. Insets show magnified regions. Scale bar, 25 μm.

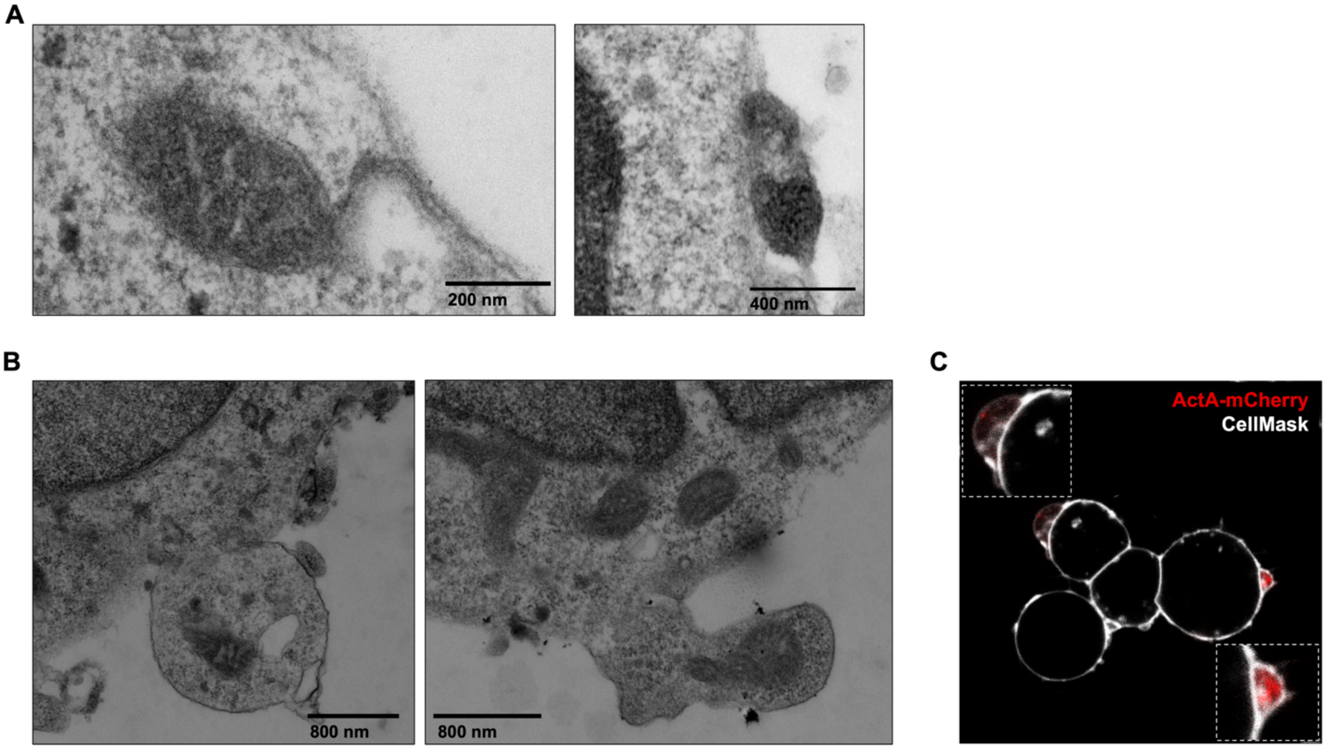


**Supplementary Fig. 8 | Ultrastructural evidence of mitochondrial membrane-associated structures. a,** Representative transmission electron micrographs of retinoblastoma cells. Representative of three biologically independent experiments. Scale bar, 200nm; 400nm. **b,** Representative transmission electron micrographs showing extracellular vesicular structures containing mitochondrial material. Scale bar, 800nm. **c,** Representative live-cell confocal images showing mitochondria-targeted ActA-mCherry within extracellular vesicular structures. Scale bar, 10 µm.

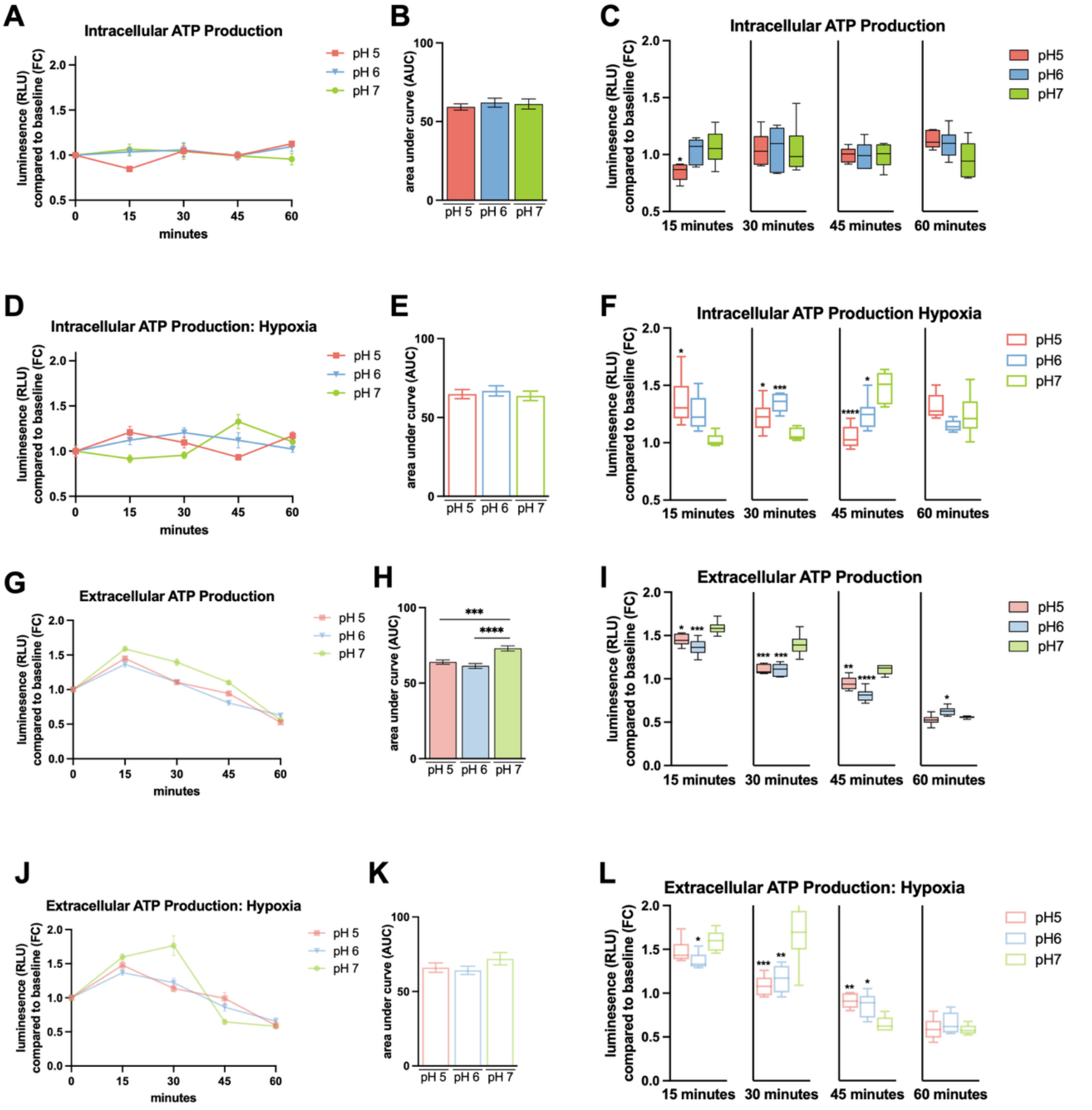


**Supplementary Fig. 9 | Extracellular acidification in non-malignant control cells. a,** Time-course quantification of intracellular ATP in mesenchymal stem cells (MSCs)cultured in 5 mM HEPES-buffered media at pH 7.4, pH 6 or pH 5 over 60 min, normalized to baseline (t = 0). Data are from three biologically independent experiments. **b,** Area under the curve (AUC) analysis of intracellular ATP under the indicated extracellular pH conditions. **c,** Comparison of intracellular ATP at 15-, 30-, 45-, and 60 min across extracellular pH conditions. Data are from three biologically independent experiments. **d-f,** As in **a-c** under hypoxic conditions (1% $O_2$). **g–l,** as in **a-f** measuring extracellular ATP production instead of intracellular ATP production. * < 0.05, *** P < 0.01, *** P < 0.001, **** P< 0.0001* versus normal media pH. Statistics**:** one-way ANOVA with Tukey's multiple comparisons test. Error bars: mean ± s.d.; box and whisker plot: min-to-max.

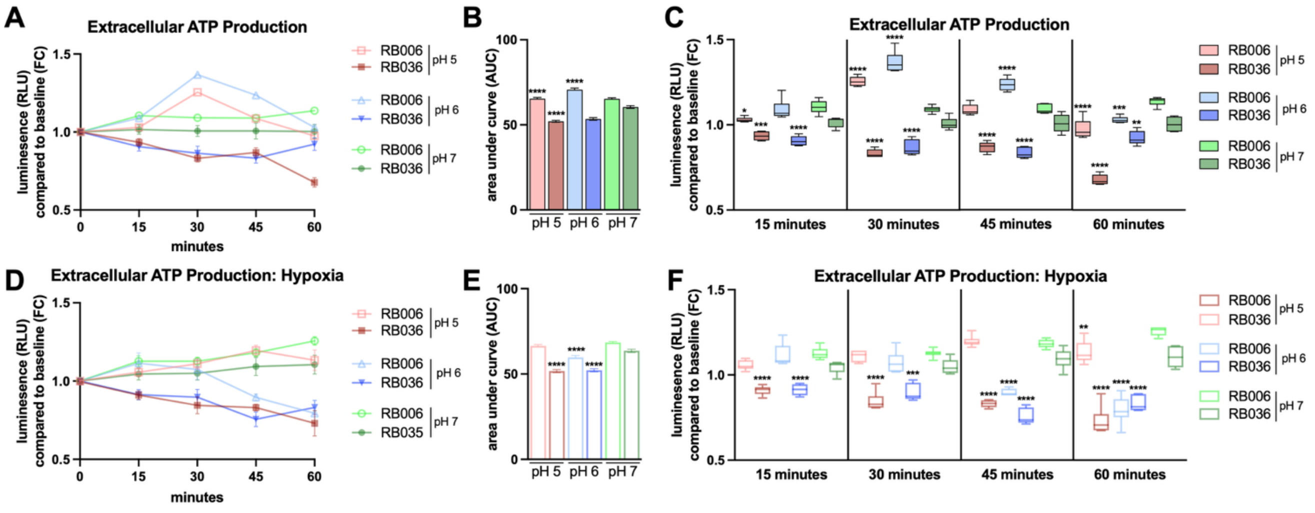


**Supplementary Fig. 10 | Extracellular ATP production in response to extracellular acidification in retinoblastoma cells. a,** Time-course quantification of extracellular ATP in RB006 and RB036 cells cultured in 5 mM HEPES-buffered media at pH 7.4, pH 6 or pH 5 over 60 min, normalized to baseline (t = 0). Data are from three biologically independent experiments. **b,** Area under the curve (AUC) analysis of extracellular ATP under the indicated extracellular pH conditions. **c,** Comparison of extracellular ATP at 15-, 30-, 45-, and 60 min across extracellular pH conditions. Data are from three biologically independent experiments. **d-f,** As in **a-c** under hypoxic conditions (1% $O_2$). * < 0.05, ** *P < 0.01,* *** *P < 0.001,* **** *P< 0.0001* versus normal media pH. Statistics**:** one-way ANOVA with Tukey's multiple comparisons test. Error bars: mean ± s.d.; box and whisker plot: min-to-max.

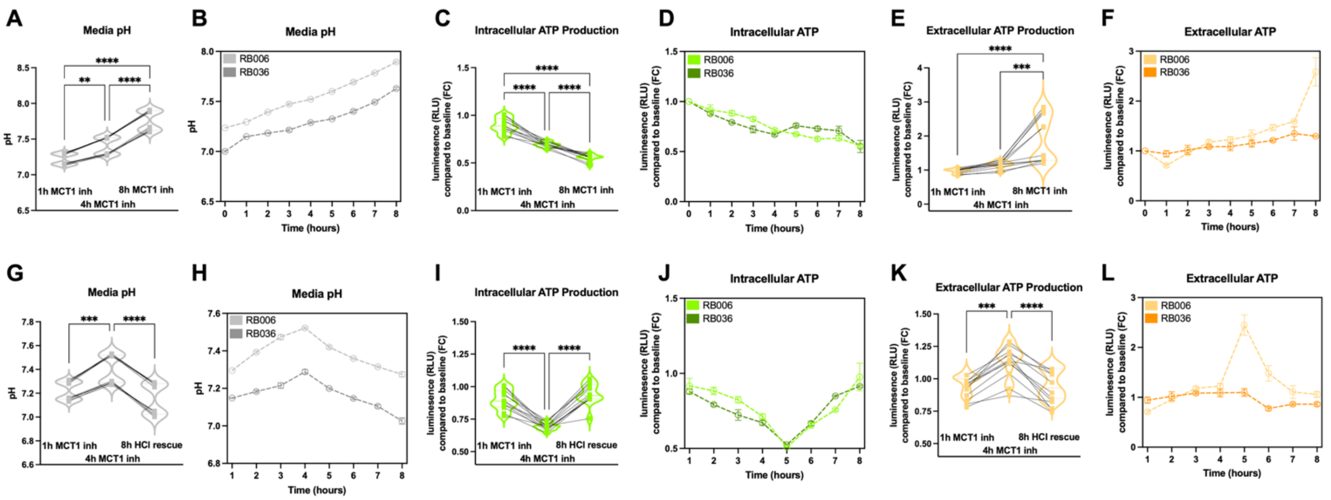


**Supplementary Fig. 11 | Intra- and extracellulr ATP production in response to MCT1 inhibition under hypoxia in retinoblastoma cells.** Cells were treated with AZD3965 (20nM) over 8 h with measurement of **a-b,** Media pH, **c-d,** Intracellular ATP production, and **e-f,** Extracellular ATP production. **g-l,** Same as in **(a-f)**; after 4 hours of AZD3965 treatment, media was re-acidified with HCl (5mM). All experiments were conducted under hypoxia (1% $O_2$). Data are from three biologically independent experiments. *** P < 0.01, *** P < 0.001, **** P< 0.0001.* Statistics**:** one-way ANOVA with Tukey's multiple comparisons test. Error bars: mean ± s.d.

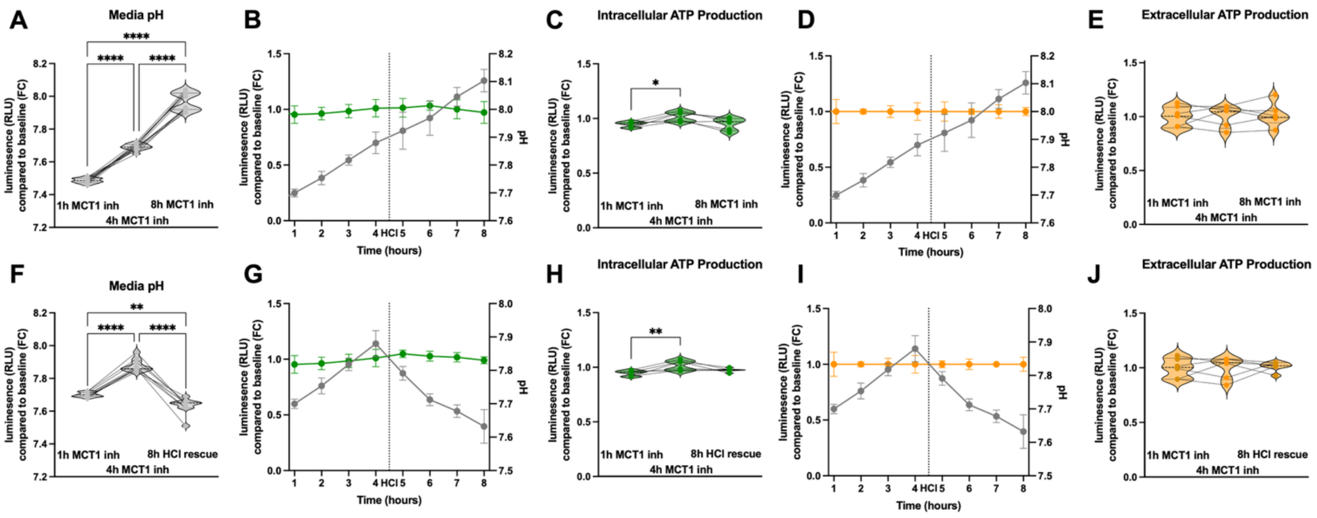


**Supplementary Fig. 12 | Control responses to MCT1 inhibition in mesenchymal stem cells. a,** Media pH in MSCs treated with AZD3965 (20 nM) over 8 h. Data are from three biologically independent experiments. **b,** Time-course of intracellular ATP (left axis) and media pH (right axis) following AZD3965 treatment. Intracellular ATP production is normalized to baseline (t = 0). Data are from three biologically independent experiments. **c,** Quantification of intracellular ATP following AZD3965 treatment at indicated time points. Data are from three biologically independent experiments. **d-e,** Same as in **(b, c)** for quantification of extracellular ATP production. **f-j,** Same as in **(a-e)**; after 4 hours of AZD3965 treatment, media was re-acidified with HCl (5mM). Data are from three biologically independent experiments. * *P <0.05, ** P < 0.01, **** P < 0.0001.* Statistics**:** one-way ANOVA with Tukey's multiple comparisons test. Error bars: mean ± s.d.

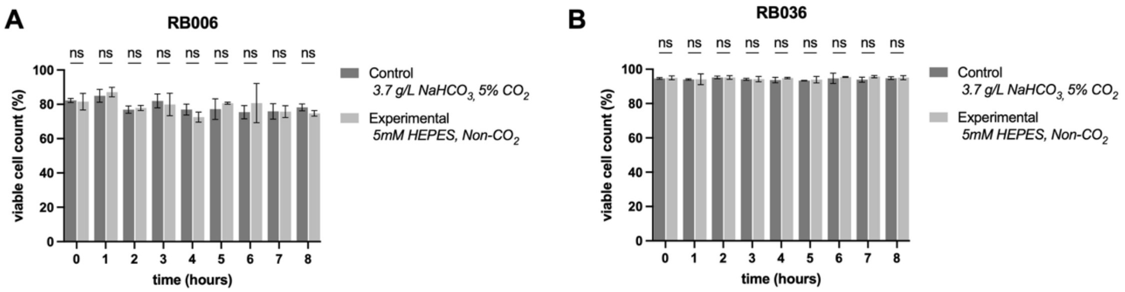


**Supplementary Fig. 13 | Control for culturing in HEPES-buffered media with non-$CO_2$ incubation. a-b,** Time-course measurement of cellular viability, measured with Trypan Blue exclusion assay, in cells cultured under normal (3.7 g/L $NaHCO_3$ w/ $CO_2$ incubation) or experimental (5mM HEPES w/ non-$CO_2$ incubation) conditions. Statistics: one-way ANOVA with Tukey's multiple comparisons test. Error bars: mean ± s.d.

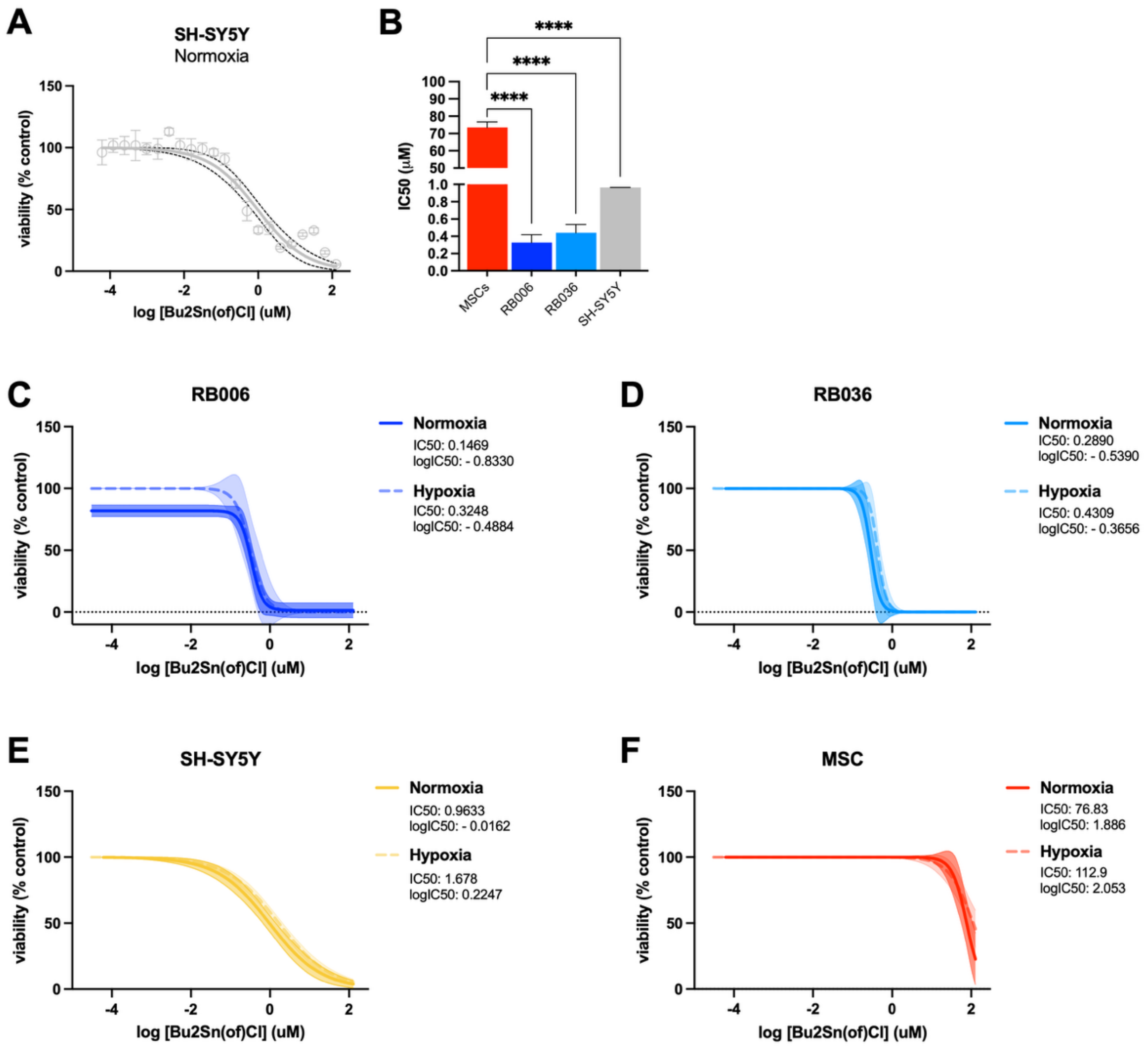


**Supplementary Fig. 14 | $F_O$ domain inhibition reduces viability in ecto-ATP synthase-expressing cancer cells. a,** Dose–response analysis of SH-SY5Y cell viability following 72 h treatment with dibutyltin-3-hydroxyflavone chloride ($Bu_2Sn(of)Cl$), quantified by CellTiter-Glo 2.0. Data are from three biologically independent experiments. Nonlinear regression was performed using a four-parameter variable-slope model. **b,** Quantification of $Bu_2Sn(of)Cl$ $IC_{50}$. **c–f,** Dose–response analysis of RB006, RB036, SH-SY5Y, and MSC viability following 72 h treatment with $Bu_2Sn(of)Cl$ under normoxia (21% $O_2$) and hypoxia (1% $O_2$), quantified by CellTiter-Glo 2.0. Data are from three biologically independent experiments. Nonlinear regression was performed using a four-parameter variable-slope model. **** *P < 0.0001*. Statistics: one-way ANOVA with Tukey's multiple comparisons test (f). Error bars: mean ± s.d.
